# The human hippocampus plays a time-limited role in retrieving autobiographical memories

**DOI:** 10.1101/2020.11.19.390526

**Authors:** Adrian W. Gilmore, Alina Quach, Sarah E. Kalinowski, Estefanía I. Gonzalez-Araya, Stephen J. Gotts, Daniel L. Schacter, Alex Martin

**Author notes:** Corresponding Author: Adrian W. Gilmore, Laboratory of Brain and Cognition, NIMH/NIH, 10 Center Drive, MSC 1366, Building 10, Room 4C104, Bethesda, MD 20892, **Email:**.

## Abstract

The necessity of the human hippocampus for remote autobiographical recall remains fiercely debated. The standard model of consolidation predicts a time-limited role for the hippocampus, but the competing multiple trace/trace transformation theories posit indefinite involvement. Lesion evidence remains inconclusive, and the inferences one can draw from fMRI have been limited by reliance on covert (silent) recall, which obscures dynamic, moment-to-moment content of retrieved memories. Here, we capitalized on advances in fMRI denoising to employ overtly spoken recall. Forty participants retrieved recent and remote memories, describing each for approximately two minutes. Details associated with each memory were identified and modeled in the fMRI timeseries data using a variant of the Autobiographical Interview procedure, and activity associated with the recall of recent and remote memories was then compared. Posterior hippocampal regions exhibited temporally-graded activity patterns (recent events > remote events), as did several regions of frontal and parietal cortex. Consistent with predictions of the standard model, recall-related hippocampal activity differed from a non-autobiographical control task only for recent, and not remote, events. Task-based connectivity between posterior hippocampal regions and others associated with mental scene construction also exhibited a temporal gradient, with greater connectivity accompanying the recall of recent events. These findings support predictions of the standard model of consolidation and demonstrate the potential benefits of overt recall in neuroimaging experiments.

## INTRODUCTION

Episodic memory refers to a collection of processes that support the retrieval of information about a spatially and temporally specific occurrence (1). A hallmark of this ability is a sense of mental time travel—of leaving the here and now and re-living or reexperiencing detailed aspects of a prior event (2). Neuropsychological and neuroimaging studies have identified a distributed collection of brain regions that support the retrieval of episodic memories, including regions within the medial temporal lobe, medial and superior prefrontal cortex, medial parietal cortex, and angular gyrus (3–5). Of particular importance seems to be the interaction of the hippocampus—a medial temporal lobe structure strongly associated with memory encoding and retrieval (6–8)—and these other regions of cortex.

Over the last several decades, a debate has arisen regarding the nature of hippocampal-neocortical interaction during the retrieval of recent, as compared to remote, events. On one hand, clinicians have long-noted a pattern by which temporally-distant memories are the best retained and the first recovered following insults such as brain injury (9, 10). Similarly, neuropsychology studies in humans (6) and lesion experiments in non-human primates (11) and rodents (12), often find temporally-graded retrograde amnesia following hippocampal damage; recently-acquired memories are lost and remote or more distant memories are spared. To account for these observations, a popular hypothesis known as Standard Model of Consolidation (SMC; 13, 14) asserts that a process of consolidation (possibly driven by hippocampal replay, 15) results in a migration of retrieval routes from the hippocampus to the neocortex, such that over time the hippocampus is no longer required for successful retrieval of a given event.

Others have argued that the critical determinate to hippocampal involvement is not time, but content and detail. The Multiple Trace and Trace Transformation (MT/TT) hypotheses—so named because they predict that each retrieval of an event is accompanied by another memory trace being formed and stored more broadly across the hippocampus—posit that the hippocampus is always required for vividly recalling memories (16, 17); memories recalled without a hippocampus are schematic and lack specific detail. Although these predictions differ substantially from those of SMC, distinguishing between the two models is unfortunately complicated by the tendency for memories to become more schematic and less detailed over time (18, 19), such that *age* and *vividness* are frequently confounded.

A number of fMRI studies in neurologically healthy participants have been conducted that speak to the question of how recent and remote events are retrieved. However, evidence has been mixed, and recent large-scale reviews have arrived at conflicting—and frankly incompatible—conclusions. For example, in a recent metaanalysis of 79 experiments, Boccia and colleagues found support for temporally-graded activity, both in the hippocampus and numerous cortical regions (5), whereas a recent review from Yonelinas and colleagues (20) concluded that neuroimaging provides “little support for the [SMC] assumption that the hippocampus becomes less involved in retrieval as episodic memories become more remote” (p. 369). In part, Yonelinas and colleagues cited the potential confounding of memory age and vividness in reaching their conclusions. In a separate recent review, Sekeres and colleagues (17) argued that neuroimaging evidence “decidedly favors the positions taken by [MT/TT]: detailed or vivid episodic memories activate the hippocampus no matter how long ago the memories were acquired.” (p. 42). Thus, over the past two years, reviews of the evidence seem to somehow simultaneously support and reject the SMC.

The debate for and against temporally graded activity in the hippocampus seems to rest largely on the subjective phenomenology of retrieved memories. From this perspective, fMRI studies of autobiographical memory have suffered in particular because of their reliance upon covert, or silent, recall. Covert recall is employed to reduce in-scanner head motion but comes at the cost of knowing the moment-to-moment content of recalled memories. In lieu of overtly recalled details, Likert-type rating scales are typically employed (either during the scan or in a post-scan session) that summarize the vividness with which a given memory was re-experienced. As a consequence, unaccounted for differences in the details associated with recent and remote memories may well obscure (or spuriously produce) effects associated with the age of a memory, contributing to the confused situation that currently characterizes the literature.

Here, we capitalized on recent advances in fMRI denoising (21) to employ *overtly spoken* in-scanner recall. Forty participants (23 female; 24.2 ± 2.8 years old) described memories from three Recall Periods: earlier on the day of scanning, a period of 6-18 months prior, and a period of 5-10 years prior. Each memory was cued using photographs of scenes and subsequently described aloud for a period of approximately two minutes (Figure 1A). A non-autobiographical control task involving a description of the same type of images was included to equate for narrative processes and possible re-encoding of details as they were being recalled and described (22), and this task served as an active baseline comparison. Memory contents were labeled using an adapted version of the Autobiographical Interview procedure (22, 23) (Figure 1B), and after synchronizing the spoken audio and fMRI timeseries (Figure 1C), contents were converted to event-related regressors to capture variance associated with every recalled detail (Figure 1D). By separately modeling details within events from the temporal distance of an event, the basic question could be asked of how the recency or remoteness of a memory, *per se*, altered retrieval-related activity. If temporally graded activity (particularly in the hippocampus) was observed, it would provide novel evidence in support of the SMC model.

**Figure 1.**
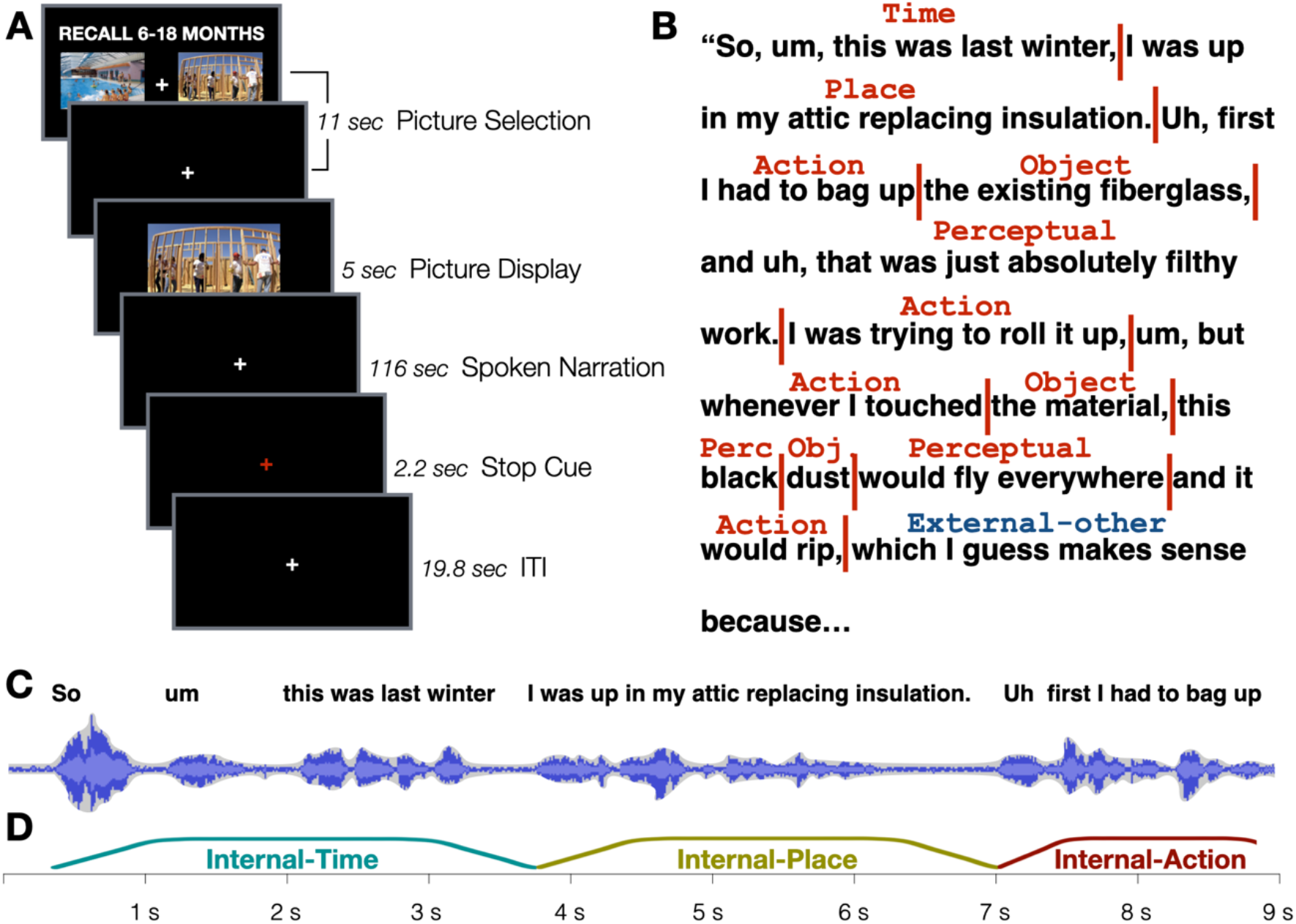
Design and analysis approach. A) Participants selected their preferred picture to use as an autobiographical retrieval cue, based on instructions at the top of the screen. Autobiographical memories were cued from 3 different recall periods (same-day, 6-18 months ago, 5-10 years ago). After viewing an enlarged version of their selected picture cue, participants described the memory as vividly as possible. A stop cue signaled the end of each trial. B) Transcripts of each description were labeled and scored for content. C) Transcript text was re-aligned with the original audio recording, and D) resynchronized with the BOLD time series so that each detail could be converted to an event-related regressor. This allowed separate modeling of event details and recall periods, the latter of which was of central interest.

## RESULTS

### Overt response scoring revealed complex event descriptions across all Recall Periods

Overt recall enabled a detailed labeling and quantification of the contents of each memory, from each condition, for each participant. Each response was scored using a variant of the Autobiographical Interview, which seeks to separate episodic and semantic contributions to memory retrieval (23). At a broad level, a given detail might be “Internal” to the event (i.e., spatially and temporally specific to the occurrence; an “episodic” detail) or it might be “External” (i.e., not specifically related to the event being described). Details were then broken down by subtype: Internal details might refer to a specific person, place, and so on, whereas External details might refer to general semantic knowledge or repetitions of previously stated information (see Table S1 for a full list of categories). Here, comparisons at both levels (Internal vs. External as well as the sub-categories within each) were compared across Recall Periods.

Average counts of Internal and External details were subjected to a repeated measures ANOVA with factors of Detail Type (Internal, External) and Recall Period (Today, 6-18 months, 5-10 years). Overall, participants were quite detailed in their responses, and a main effect of Detail Type reflected the greater number of Internal versus External details, *F*(1,39) = 106.02, *p* < .001, *η*_p_^2^ = .731 (Figure 2A). A main effect of Recall Period was also observed, reflecting temporally graded reductions in both Internal and External detail types, *F*(1,39) = 68.09, *p* < .001, *η*_p_^2^ = .193. No interaction between the factors was observed, *F*(2,78) = 2.15, *p* = .123, *η*_p_^2^ = .052. Planned comparisons of Internal detail counts indicated that a slight reduction in details was present for the most remote time period relative to the earlier conditions (6-18 months vs. 5-10 years: *t*(39) = 2.73, *p* = .009, *d* = .445; Today vs. 5-10 years: *t*(39) = 2.02, *p* = .051, *d* = .319), while an examination of External detail counts found that the most recent events contained more non-specific details than did those of later Recall Periods (Today vs. 6-18 months: *t*(39) = 2.81, *p* = .008, *d* = .445; Today vs. 5-10 years: *t*(39) = 1.92, *p* = .062, *d* = .292).

**Figure 2.**
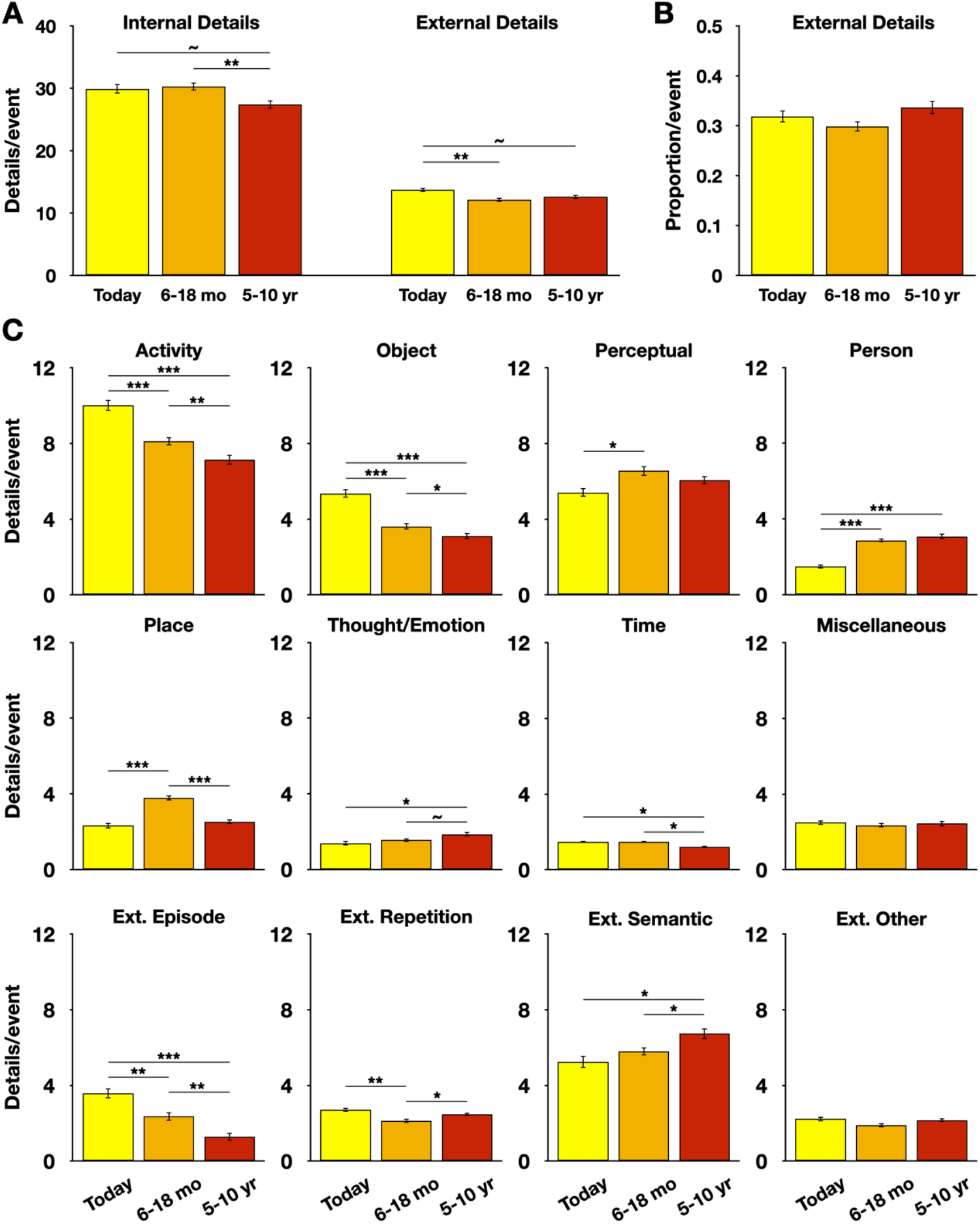
Internal and External details generated across temporal distances. A) Events from 5-10 years ago had fewer Internal details than more recent events, whereas events from the Today condition had more External details than did other Recall Periods. B) However, when taking total output into account, the overall proportion of External details did not significantly differ across Recall Periods. C) Considerable variability was present across subtypes of details, suggesting a simple “semanticizing account” of older memories may not accurately depict changes over time. Error bars denote within-subject standard error (24). * denotes *p* < .05; ** denotes *p* < .01; *** denotes *p* < .001; ~ denotes *p* < .07 (nonsignificant). For detail counts related to the Picture Description control task, see Figure S1.

The ratio of Internal-to-External details was also compared, as this may better capture broader qualitative differences across conditions. If remote memories were more schematic or semanticized in the current sample, then one would expect to see proportionally more External details as a function of Recall Period, and this may be difficult to appreciate from raw counts alone. However, no significant effect was observed, *F*(2,78) = 2.377, *p* = .10, *η*_p_^2^ = .057 (Figure 2B). Thus, although small numeric differences in Internal detail counts were present between recent and remote events, the extent to which this represented a shift in subjective memory detail over time depends on how one asks the question.

Internal and External details are composed of numerous subcategories, and the frequency of each of these categories over time was also examined. This analysis revealed a complex pattern of differences, with some categories, such as Activity and Object details, appearing more frequently in recent memories but others, such as Thought/Emotion or Person details, appearing more frequently for remote memories (Figure 2C). For still others, no effect of temporal distance was present. Thus, it was not the case that there was an across the board reduction in all types of Internal/episodic details, as one might conclude if only looking at the coarse Internal vs. External category level. Instead, these results stress that—consistent with earlier work (18, 19)—recent and remote memories differ in the composition of their overall content.

### Event descriptions from different periods did not differ in overall vividness

The prior analyses suggested that some differences in detail counts were present across Recall Periods, but their mapping to a broader change in the vividness of recall remained unclear. Better addressing this question was critical in the current experiment to rule out spurious causes of an apparent temporal gradient in neural activity (16, 17). Event descriptions were therefore deidentified and rated by independent readers using Amazon’s Mechanical Turk. In a first online experiment, participants (N = 129) were asked to rate one event description from each condition on its overall vividness on a Likert-type scale ranging from 1 (“very generic”) to 6 (“very vivid”). No differences were observed across conditions, *F*(2,256) = .779, *p* = .460, *η*_p_^2^ = .006 (Figure S2A). In a follow-up experiment, participants (N = 144) instead rated event descriptions in terms of their overall detail, with the scale ranging from 1 (“very generic”) to 6 (“very detailed”). Once again, no subjective differences were observed across conditions, *F*(2,286) = 1.78, *p* = .17, *η*_p_^2^ = .012. Both experiments were then combined into a single model; there was no effect of Experiment number, *F*(1,271) = 1.14, *p* = .286, *η*_p_^2^ = .004, no effect of Recall Period. *F*(2,542) = 1.53, *p* = .216, *η*_p_^2^ = .006, and no interaction of Experiment number and Recall Period, *F*(2,542) = 1.04, *p* = .355, *η*_p_^2^ = .004 (Figure S2B). Thus, although the types of details associated with recalled memories differed as a function of temporal distance (Figure 2), raters naïve to the hypotheses of the experiment were unable to detect any significant shift in how recent or remote events were described in the current data.

### Univariate effects of temporal distance were observed in the hippocampus, parietal cortex, and left frontal cortex

The results of the behavioral analyses highlight the shortcomings of a covert recall procedure. Without a description of how a memory is re-experienced, the activity associated with the recall of all the various Internal and External detail subcategories cannot be accounted for in an fMRI timeseries (and thus, would be aliased into regressors meant to capture activity associated with each Recall Period). Herein lies the main benefit of an overt recall paradigm: each recalled detail can be labeled and time-locked with the BOLD timeseries for each event (see Materials and Methods). This information can be converted into event-related regressors (which respect the duration and ordering of naturalistically recalled details) which can then capture transient activity associated with the recall of each identified detail (25). This separation should then provide a purer estimate of how retrieval-related activity is affected by the recency or remoteness of a memory than was previously achievable.

The mixed literature regarding temporally graded activity in the hippocampus suggested that comparing Recall Period activity within this structure could be particularly informative. Subject-specific anterior and posterior hippocampal regions—separated to reflect known functional and connectional heterogeneity across the long-axis of the hippocampus (17, 26, 27)—were interrogated for each Recall Period (Figure 3A, see Materials and Methods). Effects were tested using a repeated-measures ANOVA with factors of Subregion (Anterior, Posterior), Hemisphere (Left, Right), and Recall Period (Today, 6-18 months prior, 5-10 years prior). No main effects were observed (largest *F*-statistic: *F*(1,39) = 2.11, *p* = .128, *η*_p_^2^ = .051, obtained for Recall Period. Other *F*’s < 1). However, this must be qualified by a significant Subregion x Recall Period interaction, *F*(2,78) = 6.76, *p* = .002, *η*_p_^2^ = .148 and a marginally significant Hemisphere x Recall Period interaction, *F*(2,78) = 2.84, *p* = .065, *η*_p_^2^ = .068. The Subregion x Recall Period interaction reflects temporally graded activity differences in Posterior (Today vs. 5-10 years ago: *t*(39) = 3.05, *p* = .004, *d* = .481) but not Anterior Subregions (Today vs. 5-10 years ago: *t*(39) = .592, *p* = .557, *d* = .093) (Figure 3B). Differences took the form of greater activity for recent than for remote events. The marginal Hemisphere x Recall Period interaction reflected a smaller difference in BOLD activity between the Today and 5-10 year Recall Periods in the right hemisphere (*t*(39) = 1.02, *p* = .316, *d* = .161), as compared to left hemisphere (*t*(39) = 2.48, *p* = .018, *d* = .393). No significant three-way interaction of Recall Period, Subregion, and Hemisphere was observed, *F*(2,78) = .813, *p* = .447, *η*_p_^2^ = .020.

**Figure 3.**
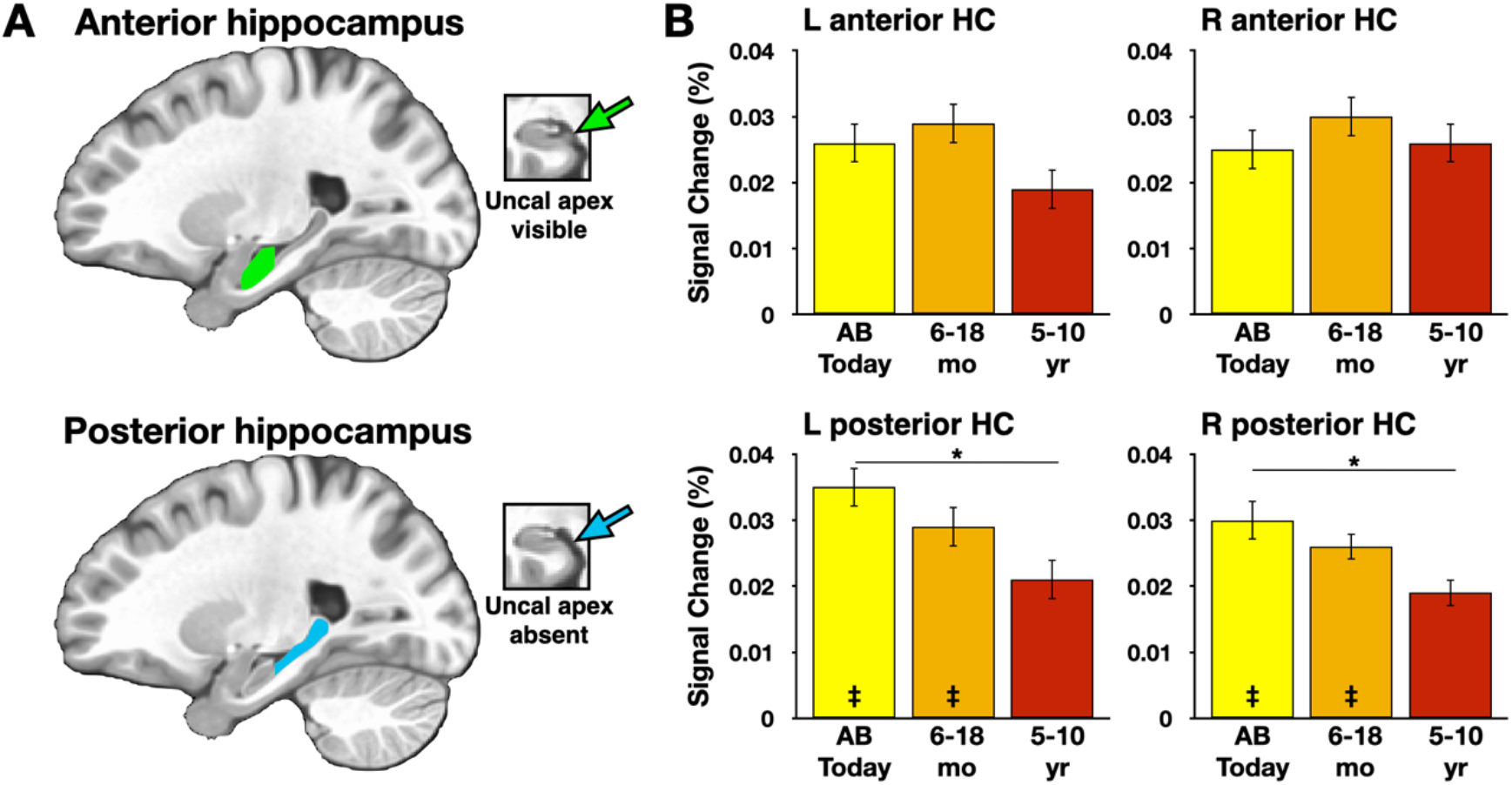
Analysis of a priori, subject-specific hippocampal ROIs. A) Anterior and posterior hippocampal ROIs were manually segmented for each hemisphere of each subject using the uncal apex as a division landmark. Inset are example anterior and posterior boundary slices. B) Posterior, but not anterior, hippocampal subregions exhibited a temporal gradient across the overt recall period, with more recent memories eliciting stronger activations than remote memories. Effects are plotted relative to the Picture Description baseline. Within posterior hippocampal ROIs, activity significantly differed from the baseline control task for the Today and 6-18 month ago, but not 5-10 year ago, conditions. Error bars denote within-subject standard error. * denotes *p* < .05; ‡ denotes significant one-sample test vs. the baseline Picture Description task, (*p* < .05, corrected for multiple comparisons); HC: hippocampus.

For the posterior hippocampal ROIs, activity in each Recall Period was subsequently compared to the active baseline task directly. The baseline task involved a verbal description of a photographic scene (equated in length to autobiographical recall durations), so basic output demands were well-matched to those of the autobiographical task. Similarly, all conditions involved the (re-)encoding of information as it was being verbally described. These considerations raise a specific question: if activity associated with recent and remote event retrieval differed, did activity also differ from the baseline task? The answer depended upon which temporal condition was being considered. Activity in both left and right posterior hippocampal ROIs differed from the baseline task for Today and 6-18 month Recall Periods (left posterior hippocampus “Today,” *t*(39) = 3.38, *p* = .012, *d* = 0.534; left posterior hippocampus “6-18 months ago” *t*(39) = 3.12, *p* = .018 *d* = 0.494; right posterior hippocampus “Today,” *t*(39) = 3.90, *p* = .002 *d* = 0.616; right posterior hippocampus “6-18 months ago” *t*(39) = 3.33, *p* = .012 *d* = 0.525; *p*-values corrected for multiple comparisons), but did not significantly differ from the baseline task in the 5-10 year ago condition (left posterior hippocampus: *t*(39) = 1.99, *p* = .318, *d* = 0.316; right posterior hippocampus: *t*(39) = 2.46, *p* = .108, *d* = .390; *p*-values corrected for multiple comparisons).

Beyond the hippocampus, effects of temporal distance were also tested at a whole-brain level. A voxelwise repeated-measures ANOVA, with the within-subject factor of Recall Period, identified regions in bilateral medial and lateral parietal cortex and the left middle frontal gyrus (Figure 4A; Table 1). Post-hoc comparisons indicated that in all of the 7 identified peaks, significantly greater activity for recent as compared to remote events was observed (Figure 4B). Notably, no regions emerged in this analysis exhibiting significantly greater activity for remote as compared to recent events.

**Figure 4.**
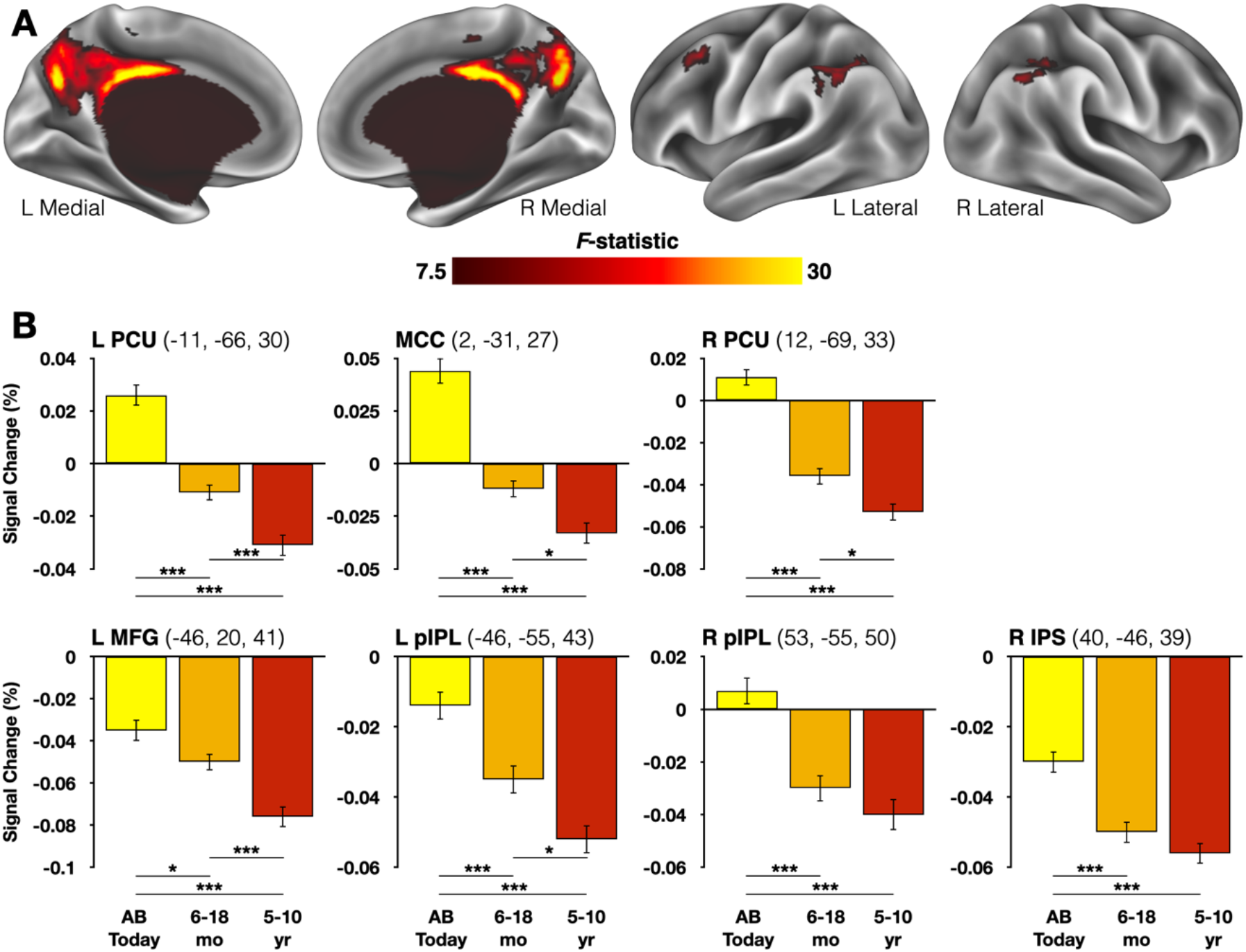
Voxelwise whole-brain analysis of temporal distance effects. A) Regions exhibiting significant main effects of temporal distance were identified in medial and lateral parietal cortex and the left superior frontal gyrus. B) Subsequent pairwise contrasts revealed that events from earlier in the same day always elicited greater activity than did more distant events, and in 7 of 7 cases a monotonic reduction in activity accompanied increasing temporal distance. Results are depicted on a partially inflated surface rendering of the human brain using Connectome Workbench software (28) and effects are plotted relative to the Picture Description baseline. Coordinates are listed in MNI152 space and refer to centers of mass for each identified region. Error bars denote within-subject standard error. * denotes *p* < .05; ** denotes *p* < .01; *** denotes *p* < .001. PCU: precuneus; MCC: mid-cingulate cortex; MFG: middle frontal gyrus; pIPL: posterior inferior parietal lobule; IPS: intraparietal sulcus.

**Table 1.** Regions identified in the voxelwise analysis of temporal distance effects. Coordinates refer to centers of mass in MNI152 space. Primary clusters were identified at *F* > 7.5; medial parietal sub-regions separated into discrete peaks at *F* > 19.8 (see Materials and Methods).

| **Region** | **X** | **Y** | **Z** | **Cluster size** | **Peak $F$-statistic** |
| --- | --- | --- | --- | --- | --- |
| Medial Parietal Cortex | -1 | -50 | 32 | 1084 | 39.19 |
| Mid-Cingulate Cortex | 2 | -31 | 27 | 113 | 39.19 |
| Left Precuneus | -11 | -66 | 30 | 43 | 38.54 |
| Right Precuneus | 12 | -69 | 33 | 38 | 34.43 |
| Left Posterior Inferior Parietal Lobule | -46 | -55 | 43 | 83 | 17.21 |
| Right Intraparietal Sulcus | 40 | -46 | 39 | 29 | 11.86 |
| Right Posterior Inferior Parietal Lobule | 53 | -59 | -43 | 29 | 13.33 |
| Left Middle Frontal Gyrus | -46 | 20 | 41 | 29 | 13.83 |

### Temporally-graded task-based connectivity was observed between regions exhibiting univariate effects and regions associated with scene construction

To complement the univariate analyses, task-based connectivity for all significant univariate ROIs (2 hippocampal, 7 neocortical) was compared for recent and remote events. These analyses took advantage of the long recall periods present in the current experiment (~2 minutes), thus providing a large number of data points for each trial. SMC would predict a temporally graded connectivity pattern that mirrors univariate effects in hippocampal regions, whereas MT/TT would predict sustained connectivity across recall periods as the hippocampus should always be involved in reactivating prior experiences. Task-based connectivity might also inform the univariate results observed in the present neocortical regions, as most of these have been associated with recognition memory performance and familiarity-related processing more than autobiographical or naturalistic recall (29–33).

In-scanner head motion is a potential problem when conducting any connectivity analysis (34). In the current case, connectivity was being compared during periods of continuous natural speech, further exacerbating potential motion concerns. It was imperative to first determine if motion differed in a measurable way across Recall Periods. This possibility was investigated in two ways. First, a repeated-measures ANOVA with the factor of average frame-to-frame motion during periods of speech found no significant effect of Recall Period, *F*(2,78) = 2.25, *p* = .112, *η*_p_^2^ = .055. Similarly, whole-brain signal variability, which is proportional to temporal signal-to-noise ratio and serves as an omnibus measure of artifactual sources of variance (35), did not differ across conditions, *F*(2,78) = 1.70, *p* = .189 *η*_p_^2^ = .042. Nevertheless, to ensure that no subtle effects related to motion might bias comparisons of connectivity across conditions, a linear mixed-effect approach was taken to compare whole-brain connectivity patterns during each Recall Period. This analysis included covariates for both the motion and signal variability measures for each trial for each participant (see also 36). As all significant effects of temporal distance in fMRI data took the form of monotonic reductions in activity over time (Figures 3,4), connectivity analyses compared correlation values specifically between the Today and 5-10 year ago conditions (see Materials and Methods).

Significant differences in connectivity between recent and remote events were observed for five of the task-based connectivity seeds. These manifested in a remarkably stereotyped pattern across the whole brain (Figure 5A, Figure S3). Consistency was quantified using a conjunction analysis approach (Table 2). The strongest overlap (≥4 maps) fell in regions of the parahippocampal cortex, the retrosplenial cortex/parieto-occipital sulcus (sometimes referred to as the “retrosplenial complex”), the posterior angular gyrus, and superior frontal cortex (Figure 5B). These regions have been strongly associated with mental time travel (and episodic simulation) in general (37), and autobiographical recall in particular (37–39). With the exception of the superior frontal gyrus, these regions are also associated with both the active visual perception of scenes and mental scene construction (40, 41), the latter of which has been argued to be a core component of vivid event recall (42–44).

**Figure 5.**
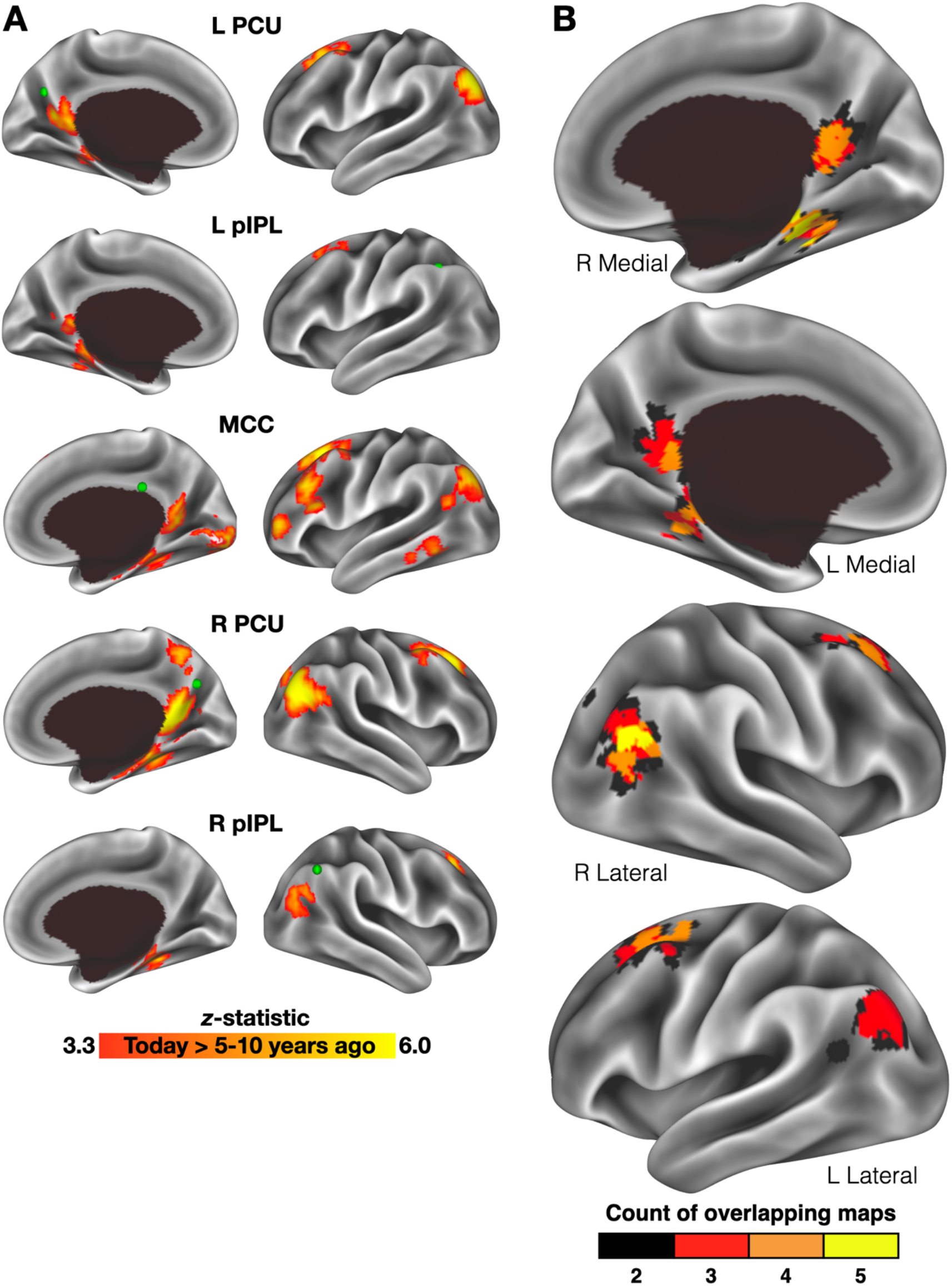
Temporally-graded connectivity was observed across seed regions. A) Regions identified in the univariate whole-brain analysis served as seeds whose connectivity was compared for recent (Today) and remote (5-10 years ago) events. Labels refer to the location of each seed region, and green nodes depict each seed’s center of mass. B) Overlap across the different seed locations was quantified by binarizing each Today > 5-10 years ago map and calculating a conjunction image. Effects were consistently observed (present in ≥4 seed maps) bilaterally in the angular gyrus, retrosplenial cortex/parieto-occipital sulcus, parahippocampal cortex, and superior frontal cortex. PCU: precuneus; MCC: mid-cingulate cortex; pIPL: posterior inferior parietal lobule.

**Table 2.** Regions identified by the conjunction analysis of temporally graded connectivity across neocortical seed regions. Coordinates refer to centers of mass in MNI152 space.

| **Region** | **X** | **Y** | **Z** | **Cluster size** |
| --- | --- | --- | --- | --- |
| *$\geq 5$ overlapping maps* |  |  |  |  |
| Right parahippocampal cortex | 28 | -34 | -17 | 9 |
| Right posterior angular gyrus | 50 | -69 | 23 | 7 |
| *$\geq 4$ overlapping maps* |  |  |  |  |
| Right posterior angular gyrus | 50 | -69 | 23 | 36 |
| Right parahippocampal cortex | 28 | -38 | -17 | 26 |
| Right retrosplenial/parieto-occipital sulcus | 8 | -55 | 8 | 25 |
| Right superior frontal cortex | 24 | 29 | 40 | 15 |
| Left retrosplenial/parieto-occipital sulcus | -8 | -49 | 4 | 13 |
| Left superior frontal cortex | -24 | 24 | 51 | 13 |
|  | -24 | 12 | 49 | 9 |
| Left parahippocampal cortex | -17 | -37 | -13 | 5 |
|  | -30 | -43 | -10 | 2 |
| Left ventral parieto-occipital sulcus | -17 | -54 | 12 | 3 |
| *$\geq 3$ overlapping maps* |  |  |  |  |
| Bilateral retrosplenial/parieto-occipital sulcus | -1 | -52 | 5 | 105 |
| Right posterior angular gyrus | 47 | -69 | 23 | 102 |
| Left posterior angular gyrus | -40 | -75 | 34 | 81 |
| Left superior frontal cortex | -24 | 21 | 48 | 80 |
| Right parahippocampal cortex | 28 | -38 | -17 | 39 |
| Right superior frontal cortex | 24 | 32 | 44 | 34 |
|  | 24 | 18 | 45 | 10 |
| Left parahippocampal cortex | -24 | -40 | -13 | 24 |
| Left middle frontal gyrus | -40 | 9 | 56 | 5 |
| *$\geq 2$ overlapping maps* |  |  |  |  |
| Bilateral retrosplenial/parieto-occipital sulcus/left parahippocampal cortex | -4 | -49 | 4 | 262 |
| Right posterior angular gyrus | 47 | -72 | 27 | 211 |
| Left superior frontal cortex | -27 | 21 | 51 | 200 |
| Left posterior angular gyrus | -40 | -75 | 30 | 138 |
| Right superior frontal cortex | 24 | 27 | 44 | 100 |
| Right parahippocampal cortex | 28 | -38 | -17 | 52 |
| Right entorhinal cortex | 21 | -21 | -18 | 12 |

The conjunction analysis results therefore led to the formulation of the final key hypothesis to be tested in this report: if autobiographical recall becomes less reliant upon the hippocampus over time, then the hippocampus should exhibit a temporally-graded pattern of connectivity with regions thought to mentally construct the scenes or contexts in which the events occur. As a means of independently identifying cortical scene-selective/scene-construction regions, a multi-category localizer dataset was collected in a separate scanning session for approximately half of the participants (N = 22, see Materials and Methods). This procedure involved the visual presentation of 9 different stimulus categories, and a contrast of activity during blocks of scenes against all other categories identified a group-level mask of scene-selective cortex, which includes—and extends beyond—the regions consistently observed in the conjunction analysis (41) (Figure 6A). The timeseries of all voxels within the scene-selective mask was correlated with each posterior hippocampal seed region and averaged in a trialwise fashion. As before, a linear mixed effect modeling approach was taken, with motion and signal variability covariates associated with each specific trial in each condition for each participant.

**Figure 6.**
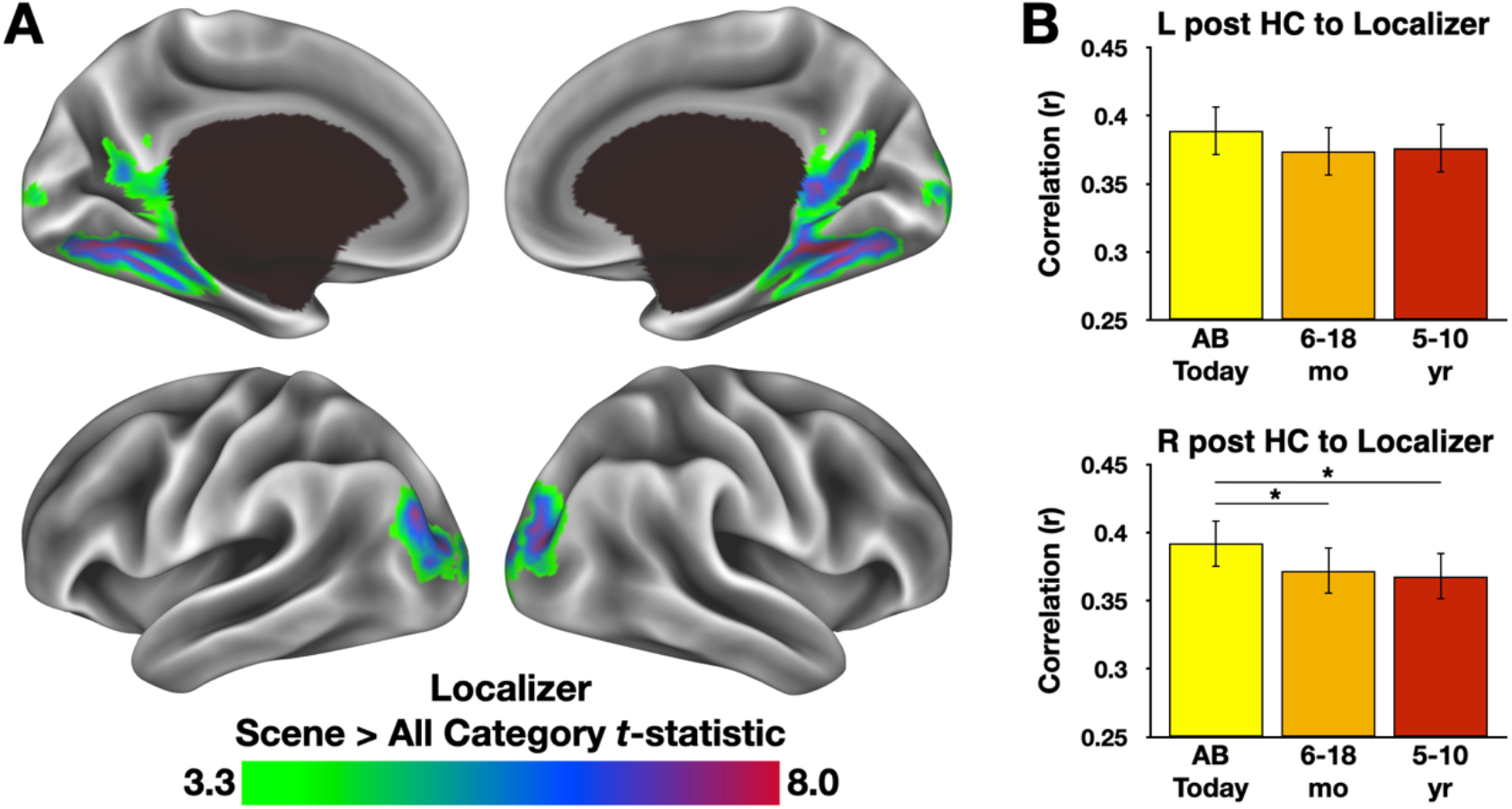
Scene-selective cortex exhibits a temporal gradient in task-based connectivity to the posterior hippocampus. A) A contrast of Scene and other category blocks in a multi-category localizer enabled the identification of scene-selective cortical regions. B) Comparing the average correlation of scene-selective voxels with posterior hippocampal regions across Recall Periods revealed a significant difference in task-based connectivity in the right posterior hippocampus and a non-significant tendency in the left posterior hippocampus. Error bars reflect standard error of the mean. * denotes *p* < .05.

The contrast of connectivity strength for recent and remote events revealed a significant difference for the Today vs. 5-10 year ago Recall Periods in the right posterior hippocampus, *t*(611) = 2.51, *p* = .012, as well as a significant difference between the Today and 6-18 month ago conditions, *t*(611) = 2.13, *p* = .033. A similar but non-significant tendency was observed in the left posterior hippocampus for the Today vs. 5-10 year ago Recall Periods, t(611) = 1.43, *p* = .154 (Fig. 6B). Thus, just as univariate effects within the hippocampus were identifiable following a targeted analysis, so too were temporally graded hippocampal-cortical interactions, with the latter occurring in regions of cortex thought to process the mental construction of scenes. Importantly, the observed pattern does not appear explainable by univariate BOLD activity difference across voxels the mask, as no significant differences in activity were observed between the Today and 5-10 year ago Recall periods, *t*(39) = 0.26, *p* = .795, *d* = .042.

## DISCUSSION

The work in this report used spoken in-scanner recall to investigate the nature of recent and remote memory retrieval. After identifying the moment-to-moment content of recalled memories using an established scoring approach (22, 23), we observed temporally-graded activity emerge in posterior hippocampal ROIs as well as in a cortical network often associated with the processing of stimulus familiarity. Regions exhibiting temporal gradients also showed a graded correlation during recall with regions thought to support the mental construction of scenes. Taken as a whole, the results of this study help to clarify an ambiguous literature, lending support to the SMC hypothesis, and emphasizing the utility of overt recall in the cognitive neuroscientific study of autobiographical memory.

### Overt recall provides new insights into an old problem

The subjective experience of mental time travel is a critical aspect of an episodic memory (1). To date, fMRI studies of autobiographical recall—even those that have manipulated the remoteness or recency of recalled events — have relied upon covert retrieval, typically with something like a Likert-type rating scale indicating the subjective vividness or richness of the recalled memory. Such ratings can be efficiently collected but necessarily provide an impoverished view into the rich phenomenology of episodic recall. In contrast, the current approach goes well beyond this simple rating scale in that it uses overt, recordable descriptions to assess phenomenology. This approach does not, of course, provide a direct window into the minds of participants—such a capability remains sorely awaited by psychologists and neuroscientists alike—yet it provides a means of accounting for, at least to some degree, dynamic effects associated with the recollection of different event details (25). By accounting for such activity, the current approach provides a means of separating the content of memories from their temporal distance in a manner not previously possible, and certainly in a manner that exceeds the variance that can be accounted for using trial-level Likert rating. Similarly, the current approach also extends beyond what was possible based on previous back-sorting approaches. For instance, Spiers and Maguire (45) collected detailed post-scan narration following a memory-guided navigation task, and used these narratives to code certain behaviors at decision points along the route. This approach was remarkably clever, yet it restricted time-locking only to specific decision points and thus could not account for the rich variety of thoughts that accompany the recall of a past episode.

### Temporal distance effects were specific to posterior hippocampal regions

Having labeled, quantified, and modeled the dynamic moment-to-moment content of recalled experiences (complete with the tangents, meta-cognitive statements, and nonlinear descriptions associated with naturalistic recall), and after testing for—but failing to observe—broader qualitative differences across event descriptions, we could then ask how recent and remote events might be differently retrieved at the neural level. Within the hippocampus, posterior but not anterior regions exhibited significant temporal distance effects across participants. Although the SMC does not distinguish between anterior and posterior hippocampal subregions in its predictions, the directionality of differences is consistent with the SMC hypothesis. Furthermore, the current data echo recent meta-analytic findings of posterior (not anterior) hippocampal regions exhibiting temporally-graded activity during autobiographical recall (5).

The behavioral results obtained through overt recall appear to rule out an alternative “semanticizing” account that might otherwise explain the fMRI results (i.e., the changes in memory content are appreciably more complicated than a general reduction in Internal details; Figure 2, Figure S2). Although this may appear surprising given the strong concerns raised by proponents of the MT/TT framework, the current pattern is not without precedent, and relevant prior fMRI studies have also reported a similar number of total details for recent and remote events (e.g., 46, 47). The most straightforward explanation is therefore that there is at least some effect of temporal distance on hippocampal activity during the recall of autobiographical events, and that this effect is consistent with predictions made by Squire and others over the past several decades in the form of the SMC framework (13, 14).

### On the importance of an active control condition in testing for a temporal gradient

A challenge in studying hippocampal contributions to retrieval is that it is always actively encoding one’s ongoing experiences (48), including information that is recalled during retrieval (49). Indeed, such re-encoding is central to the MT-TT framework (16, 17). A natural problem thus becomes the separation of activity related to retrieving event details from the re-encoding of the same details. Importantly, the nature of the control task used in this experiment—and the results of comparisons between the control and Autobiographical recall tasks—suggests that re-encoding is unlikely to explain the current findings. As has been noted in prior behavioral work that used the same control task (22, 50), visual processing of picture cues and required verbal outputs are well-matched across the Autobiographical Recall and Picture Description control conditions. In both cases, these aspects of each trial need to be (re-)encoded (see also 51 for related reasoning using a conceptually similar control). However, what is necessarily absent from the Picture Description condition is an existing episodic memory representation. Differences in activity between the baseline and Autobiographical Recall conditions should therefore be attributable to retrieval-related processing. Within posterior hippocampal regions, activity differed from this encoding baseline in 2 of the 3 Recall Periods (Figure 3B)—those associated with the more recent conditions. It was only in the most temporally remote condition that significant differences between the Picture Description and Autobiographical Recall conditions were not observed. The obtained results once again seem to favor predictions of the SMC hypothesis.

### Cortical signatures of temporal distance during overt recall

Beyond effects within the hippocampus, temporal distance effects were also observed in a small collection of neocortical regions. Effects manifested in the form of greater activity accompanying recent, rather than remote memories; a pattern not cleanly predicted by either the SMC or MT/TT models. SMC would predict either no change or greater univariate activity for remote events (complementing reduced activity in/reliance on the hippocampus), whereas MT/TT predicts that activity should be based on retrieved event features rather than temporal distance. However, the identified cortical regions have all been associated with recognition memory (cf. 52), and those in parietal cortex align particularly well with the parietal memory network, which is thought to support the recognition of and orienting toward familiar stimuli (32, 53, 54). If one assumes that objective recency is accompanied by a strong subjective familiarity when recalling an event, then the current observations coincide well with prior work, both with studies involving externally-presented stimuli (e.g., 53, 54, 55) as well as those requiring repeatedly imagining complex events (e.g., 56, 57, 58). Furthermore, the foundation of parietal memory network knowledge consists of laboratory studies that typically involve very short (minutes-long) delays between subsequent exposures. It is an intriguing possibility that this same network may support a sense of familiarity for complex events that occurred hours, months or even years ago, and follow-up work may offer a mechanistic explanation for a means by which temporal distance estimates are made when thinking back to past episodes.

There is evidence of univariate (59) or multivariate (60, 61) effects of temporal distance in ventromedial prefrontal cortex (vmPFC)—sometimes manifesting as nonmonotonic shifts in activity over time (62). However, temporal distance effects were not reliably observed in recent meta-analyses (5), nor were any observed in this report. Thus, while it seems clear that vmPFC is important for autobiographical memory recall in general (e.g., 3, 30, 63, 64), the precise conditions under which it exhibits a temporally graded activity pattern require further clarification.

### Interactivity between the hippocampus and scene construction regions during recall

It is thought that scene construction is a core process underlying episodic autobiographical recall (42–44). Indeed, a memory without a spatial context fails to meet the basic criteria of an episodic memory (1), and lesions to regions associated with scene construction reliably produce memory impairments and amnesia (65). Given the centrality of scene construction to autobiographical recall, one might expect to see a similar engagement of regions that support scene construction across temporal distance, assuming detailed recollections in each case. Consistent with this expectation, BOLD activity within scene-construction regions did not significantly differ between the most recent and most remote conditions. However, the data also demonstrate that connectivity between scene-construction regions (localized independently within a subset of the overall sample) and the posterior hippocampus is reduced in a temporally graded fashion. In other words, although the scene construction regions themselves seem to be similarly engaged when recalling recent or remote events, the nature of the interactions between scene construction regions and the posterior hippocampus differs. Of relevance here, the differences observed are consistent with a reduced hippocampal engagement over time, as predicted by the SMC model.

### Conclusion

In sum, the present data reveal that overt, in-scanner recall provides evidence that supports predictions made by the SMC hypothesis: the hippocampus exhibits temporally-graded reductions in activity during autobiographical recall, as well as reduced hippocampal-cortical interactivity with regions thought to serve a core aspect of autobiographical memory. At the same time, regions thought to be important for the recognition of familiar stimuli also appear sensitive to the recency or remoteness of memories that occurred months or even years ago, suggesting an important phenomenological role for these regions in retrieval processes in general. Finally, these data encourage the utilization of overt in-scanner recall to study human cognition and demonstrate that technical concerns related to motion need not hold researchers back from collecting larger and richer datasets.

## MATERIALS AND METHODS

### fMRI Participants

46 participants were recruited from the NIH community and greater DC metro area. Of these, 3 were excluded for excessive movement (with 5 or more of the 8 task scan runs excluded based on motion or were missing all of one type of task scan, see below), 1 was excluded due to technical problems encountered while scanning, and response data were lost from an additional 2 participants. The remaining 40 participants (23 female) had a mean age of 24.2 (SD ±2.8 years, range 20 to 34), were right-handed, had normal or corrected-to-normal vision, were native speakers of English, and were neurologically healthy. Informed consent was obtained from all participants prior to their participation, and the experiment was approved by the NIH institutional review board (clinical trials number NCT00001360). All participants were monetarily compensated for their time.

### Online (Mechanical Turk) Participants

In Online Experiment 1, 210 participants over 18 years of age and who were located in the US completed the task. Data from 81 participants were excluded for failing to comply with task instructions (see Online Experiment 1 Procedure). The retained 129 participants (50 female, 2 of unreported gender) ranged from 18-69 years old (mean = 35.8± 12.1 years; 5 participants did not disclose their exact age).

Online Experiment 2 followed the same participant requirements and exclusion criteria as Online Experiment 1. 144 of 199 participants were retained (60 female, 7 of unreported gender). Ages ranged from 19-72 years (mean = 36.6± 10.5 years; 4 participants did not disclose their age). Workers who completed Online Experiment 1 could not participate in Online Experiment 2.

For both Online Experiments, informed consent was obtained from all participants prior to their participation. The experiments were approved by the NIH institutional review board (clinical trials number NCT00001360), and all participants were monetarily compensated for their time.

### fMRI Stimuli

Autobiographical and Picture Description stimuli consisted of 48 pictures depicting people in different locations undertaking various activities (e.g., ordering at a café). Some of these images have been described previously (22, 50); the remainder were newly acquired for this study via internet search. Images were presented in color against a black background and were sized at 525 x 395 pixels.

Multi-category localizer task stimuli consisted of 120 images of 8 different categories (abstract shapes, animals, body parts, static dots, faces, non-manipulable objects, scenes, phase-scrambled images, tools, and words), as well as 5 images that cued different movements, from a larger collection described previously by Stevens, Tessler, Peng and Martin (66). Images were 600 x 600 pixels and were gray scaled.

Stimuli for all tasks were presented using PsychoPy2 software (67) (RRID: SCR_006571) on an HP desktop computer running Windows 10 (display resolution: 1920 x 1080 pixels).

### Online Experiment stimuli

Stimuli used in both Online Experiments were modified verbal reports from the main fMRI experiment. 210 de-identified event descriptions were selected from each Recall Period. Event triads were then formed by randomly selecting (without replacement) one event from each Recall Period. Event descriptions were presented in black, 12-point Arial type against a white background.

### Autobiographical Recall task

In this task, participants overtly retrieved autobiographical memories of different ages in response to picture cues. Trials began with a screen that instructed participants to think back to one of three different recall periods (earlier in the same day, 6-18 months ago, 5-10 years ago), and provided participants with two different picture cues (Figure 1). Participants had 11 s to select (via button press) the picture they preferred to use as an autobiographical memory cue. Participants were provided a choice of images to reduce instances of event recall failure. Image pairings were shuffled across participants and rotated across conditions, with a subset of more typical scenes reserved for the “Today” condition. Following their selection, the images were removed until the end of the 11 s selection period, at which point an enlarged version of selected image was presented in the center of the screen for a 5 s period. Participants were instructed during this time to use the picture to help remember an event from the cued time period.

Following the 5 s picture presentation, the image was removed and replaced with a white crosshair for 116 s. During this time, participants were instructed to narrate the cued memory with as much detail as possible for the full duration of the trial. Participants were instructed that events should be unique (i.e., the same event should not be described multiple times) and should be specific in time and place (i.e., should reflect unique episodes rather than routine recurrences). In cases where participants ceased early in the trial (e.g., with ≥ 20 s remaining), they were given a general prompt by the experimenter. This took the form of the question “Are there any other details that come to mind?” as described by Levine, Svoboda, Hay, Winocur and Moscovitch (23). Participants heard the question via their noise-cancelling headphones. Such prompts were rare (averaging <1 occurrence per individual). At the end of the narration period, the white fixation cross changed to a red color for 2.2 s, which signaled the end of the trial. Trials were separated by 19.8 s of fixation, and three trials (one per Recall Period) were included per scan run. Six autobiographical task runs were collected for each participant, resulting in 18 total Autobiographical Recall trials. The order of Recall Periods was counterbalanced across runs and participants.

### Picture Description task

This control task required descriptions of complex photographs and was modeled after Autobiographical Recall trials. Participants were cued to describe what was occurring in a given picture, and the instruction was accompanied by two images. Participants had 11 s to select (via button press) their preferred picture. Image pairings were shuffled across participants. The images were removed until the end of the 11 s selection period, when an enlarged version of the selected image was presented for 5 s. Participants were instructed to scrutinize the image during this time. The image was then replaced with a white crosshair for 116 s. During this time, participants were instructed to describe the image with as much detail as possible for the full duration of the trial. The white fixation cross changed to a red color for 2.2 s at the end of each trial. As in the Autobiographical Recall task, participants were prompted if they ceased speaking early in a trial. Trials were separated by 19.8 s of fixation, and three picture description trials were included per scan run. Two runs of the Picture Description task were collected for each participant (for 6 total Picture Description trials) and were interleaved with the Autobiographical Recall scan runs. Ordering was counterbalanced such that an equal number of Autobiographical Recall runs preceded or followed the Picture Description scans.

Prior to scanning, participants were instructed as to the nature of the experimental tasks. They practiced engaging in both task types and were given additional instruction/practice cycles if autobiographical events were not initially specific in time and place or otherwise episodic in nature.

### Multi-category localizer task

In a separate session, approximately half of the participants completed a 1-back task meant to serve as a multi-category functional localizer. Participants were presented with blocks of images from each included category and were directed to press a button when they noticed a repetition of the same image. 20 images were presented in each block, and each image was presented for 300 ms followed by 800 ms of fixation. Blocks were separated by an 11 s period of fixation. All blocks within each run contained different types of stimuli and were placed pseudorandomly. 1 block per run served as a “motor” localizer in which visual cues directed participants to flex their left or right fingers, left or right toes, or flick their tongue against their front teeth. This motor localizer was based on that used in the Midnight Scan Club dataset (68). Six runs of the multi-category localizer were collected for each participant.

### Audio recording and in-scanner speech

Participants spoke their descriptions into an Optoacoustics FOMRI-III NC MR-compatible microphone with active noise cancellation. Output from the microphone was passed into an M-Audio FastTrack Ultra 8-R USB audio/MIDI interface (inMusic, Cumberland, RI) and was recorded using Adobe Audition on a Dell Precision M4400 laptop. Spoken audio tracks were transcribed for subsequent text analyses (see below). A secondary track captured a square wave pulse set at the onset of each stimulus presentation to allow precise syncing of audio tracks and scan onset times.

Prior to beginning the experiment, participants were given a chance to practice speaking while the scanner was running. Real-time estimated motion traces were examined by the experimenter (AG), who provided feedback to participants over the MR-compatible and noise-cancelling headset regarding the severity and types of motion that were being observed. Motion estimates were generated using a realtime AFNI implementation and included 3 translation and 3 rotation parameters for each TR immediately after it was collected.

### Alignment of spoken responses to BOLD timeseries data

Spoken response tracks were processed in Audacity 2.3 (audacityteam.org) to reduce background noise. All spoken responses were transcribed and were then checked against the original recorded audio to ensure that they were free of typographical errors. A python-based text-to-speech alignment tool from the University of Pennsylvania Department of Linguistics, (p2fa; 69), was used to synchronize the text and spoken audio for each event; outputs were manually edited to correct misalignments using Praat (70) v6.0.48 (RRID: SCR_016564). Finally, the speech onset response time (RT) for each event was calculated by comparing the time difference between the onset of the picture cue (recorded in a secondary audio track as a square pulse) and the onset of speech. As p2fa temporally aligned all words with the initial word, onset times and durations could be calculated for every spoken word and phrase. Timing information was used for modeling univariate retrieval effects (explained in more detail below).

### Transcript scoring

Transcribed autobiographical memories and picture descriptions were scored using an adapted version of the Autobiographical Interview scoring system (23), modified to accommodate both memory and picture descriptions in a manner similar to that reported by Gaesser, Sacchetti, Addis and Schacter (22), with additional modifications as described below. Briefly, this approach segments different details provided by participants as either being “Internal” (i.e., episodic details related to the specific event being described) or “External” (i.e., details that were from unrelated episodes, were semantic/non-specific in nature, or were repetitions of previously described details). For picture description trials, any details that depicted elements in the picture were considered part of the central event. For autobiographical recall trials, the coder identified the “central” event for purposes of scoring if multiple events were described. One notable update to the scoring procedure concerns the use of the “Event” details category. As originally conceptualized, these refer to a broad range of details including persons present, actions/reactions, weather conditions, and “happenings”. However, as it is known that different cortical regions support the processing of different concepts and object properties (71), the “Event details” category was broken down into more specific detail types (i.e., people, objects, activities, miscellaneous; see 25). A full list of detail types is presented in Table S1.

Detail scoring was conducted by 3 separate coders, each of whom scored a subset of the overall participants. Interrater reliability was assessed on the basis of pilot data via interclass correlation analysis using a 2-way random model. Training and subsequent testing resulted in a strong overall reliability across Internal and External detail categories, ICC(2,3) = 0.92.

Across the 3 Recall Periods, the overall frequency of Internal and External detail types, as well as the subtypes within each broad category, were compared using repeated measures ANOVAs and follow-up pairwise comparisons (two-tailed). Given the large number of ANOVAs conducted in investigating this aspect of the data, an FDR approach (72) was selected for multiple comparison correction, requiring *q* < .05.

### fMRI data acquisition

Scanning was performed using a General Electric Discovery MR750 3.0T scanner, with a 32-channel head coil. Functional images were acquired using a BOLD-contrast sensitive multi-echo echo-planar sequence (Array Spatial Sensitivity Encoding Technique [ASSET] acceleration factor = 2, TEs = 12.5, 27.7, and 42.9 ms, TR = 2200 ms, flip angle = 75°, 64 x 64 matrix, in-plane resolution = 3.2 x 3.2 mm). Whole-brain EPI volumes (MR frames) of 33 interleaved, 3.5-mm-thick oblique slices were obtained every 2.2 s. Slices were manually aligned to the AC-PC axis. A high-resolution T1 structural image was also obtained for each subject (TE = 3.47 ms, TR = 2.53 s, TI = 900 ms, flip angle = 7°, 172 slices of 1 x 1 x 1 mm voxels).

Foam pillows helped stabilize head position for all participants, scanner noise was attenuated using foam ear plugs and a noise-cancelling headset. The headset was equipped with speakers that facilitated communication with the subject during their time in the scanner. A sensor was placed on each participant’s left middle finger to record heart rate, and a respiration belt monitored breathing for each subject.

### fMRI preprocessing

fMRI data were preprocessed using AFNI (73) (RRID: SCR_005927) to reduce noise and facilitate across-subject comparisons. Initial steps included frame-by-frame rigid-body realignment to the first volume of each run (3dvolreg), slice-timing correction (3dTshift), and despiking to remove large transients in the timeseries (3dDespike). The first 4 frames of each run were discarded to remove potential T1 equilibration effects. Following these initial steps, data from the three echoes acquired for each run was used to remove additional noise using multi-echo independent components analysis (ME-ICA) (21, 74, 75). Briefly, this procedure calculates a weighted-averaging of the different echo times to reduce thermal noise within each voxel, and then uses spatial ICA and the known properties of T_2_^*^ signal decay to separate putatively neuronal components from those thought to be artefactual in nature, including thermal noise and localized effects of head motion (76). Components are retained if they show both good fit with a model that assumes a temporal dependence of signal intensity and a poor fit with a model that assumes temporal independence of signal intensity (74). Components were selected using the default settings of AFNI’s *tedana.py* function. Following ME-ICA processing, data from each scan run were aligned across runs and registered to each individual’s T1 image. Data from each participant were then resampled into 3-mm isotropic voxels and linearly transformed into standardized atlas space (TT_N27).

### GLM-based fMRI data analysis

All task scans (6 Autobiographical Recall, 2 Picture Description) consisted of 210 MR frames (214 prior to initial frame discarding) and lasted 7 minutes, 51 s in duration. Run-level motion summaries reflected the average of first differences in frame-to-frame head position across each scan run and were obtained using the AFNI program @1dDiffMag. Any runs exceeding 0.2 mm/TR were excluded. This resulted in the exclusion of three participants (two of whom had 7/8 runs excluded, and a third had 5/8 runs excluded, including both picture description runs). Additionally, 2 autobiographical task runs were excluded from 4 additional participants based on this motion criterion, and 1 autobiographical task run was excluded from 5 participants. In the remaining task runs, average motion was 0.13 mm/TR for the Autobiographical Recall task and 0.12 mm/TR for the Picture Description task, a modest yet significant difference, *t*(39) = 3.34, *p* = 0.002.

Functional data from each subject were smoothed using a 3 mm FWHM Gaussian kernel to account for intersubject anatomical variability, normalized by the mean signal intensity of each voxel prior to analysis, and linearly detrended to account for scanner drift effects. Analysis was conducted using a general linear model (GLM) approach via AFNI’s 3dDeconvolve function. Data from each time point were treated as the sum of all effects present at that time point. The initial picture selection period was modeled using a single HRF across all trial types convolved with a boxcar of 11 s duration (implemented using AFNI’s ‘BLOCK’ function). The subsequent Picture Display period was also modeled with a single HRF convolved with a boxcar of 5 s duration. The analysis of recall effects utilized a mixed block/event related design (77–79). Separate regressors modeled sustained effects related to the narration periods of the 3 Autobiographical Recall conditions as well as the Picture Description narration period. These convolved an HRF with a boxcar of 118.2 s duration in all cases. Additional regressors coded for transient effects associated with the type of detail being described throughout each narration period, as detailed in “Transcript Scoring” above. This added an additional 12 columns to the design matrix and provided a means to account for transient, detail-related reactivation effects that differed throughout and across each described event (reported and discussed by 25). Reactivation-related effects for each detail type were modeled using AFNI’s ‘dmblock’ response function so that the durations of each described detail could be included in the GLM; multiple consecutive details of the same type were modeled as a single event. By modeling transient effects, estimations of sustained effects should more accurately reflect basic differences associated with the temporal distance of an event. Finally, 6 estimated motion parameters (3 translational, 3 rotational) were included as regressors of non-interest.

### Hippocampal ROI definition

Of specific interest were effects in the hippocampus that could be associated with different task conditions. Subject-specific hippocampal masks were generated with Freesurfer (v6.0; RRID:SCR_001847), and each mask was manually segmented into anterior and posterior sub-regions using the uncal apex as a landmark of separation (26). Masks for each region for each subject were then resampled to the resolution of the EPI data.

### Calculating effects of temporal distance

Of primary interest was BOLD activity during the different Autobiographical Recall task conditions. Activity was averaged for each recall condition across all voxels in each hippocampal region of interest (ROI) in each subject’s native space, with the picture description task serving as a baseline comparison condition. A repeated measures ANOVA with a single factor of Recall Period (3 levels: Today, 6-18 months ago, 5-10 years ago) was conducted for each of the 4 hippocampal subregions. Where significant, ANOVAs were followed up with paired samples, two-tailed *t*-tests to understand the nature of the differences. One-sample *t*-tests were subsequently conducted in posterior hippocampal ROIs to compare the significance of each response versus the baseline Picture Description condition (requiring FWE *p* < .05).

The same analysis was repeated, this time at the voxelwise whole brain level. The resulting statistical map was corrected to a whole-brain *p* < .05 by requiring a voxelwise significance of *p* < .001 (*F* > 7.5) with a minimum cluster extent of 22 voxels. These values were calculated using AFNI’s 3dClustSim and its updated non-Gaussian autocorrelation function (80, 81), which was released following the findings of Eklund, Nichols and Knutsson (82). In the initial voxelwise analysis, one large midline cluster (*k* = 1084 voxels) was identified at the nominal voxelwise correction level with 3 clear and distinct local maxima present within it. These local maxima within the cluster were separated by incrementing the minimum voxelwise *F*-statistic in steps of 0.1 until all were separated; this was achieved at *F* ≥ 19.8 (*p* ≤ 1.1 x 10^-7^) and these separated clusters were used to define bilateral precuneus (PCU) and mid-cingulate cortex (MCC) regions.

As with the hippocampal ROI analysis, pairwise comparisons of activity associated with each Recall Period were then conducted for each identified cluster to better interpret the directionality of observed effects.

### Task-based connectivity analysis

To better understand the observed univariate effects in the current dataset, task-based connectivity was compared across the different Autobiographical Recall periods. For each subject, 53-TR (116.6 s) windows for each event were notched out of the residual timeseries (i.e., with conditional effects regressed out as described previously), beginning the TR after description period began and ending at the stop-cue for each trial. This corresponds to the time windows interrogated when estimating motion and signal variability, which were included as trial-level quantitative covariates to help ensure that observed correlations were not driven by nuisance effects (for similar reasoning, see 36).

The 2 posterior hippocampal ROIs, as well as the regions identified in the univariate temporal distance analysis, served as seed regions in a whole-brain analysis. The timecourse of activity for each seed for each trial was correlated with all voxels in the brain for each trial of each condition. The resulting correlation maps were Fisher transformed and entered into a linear mixed-effects model using AFNI’s 3dLME (83), with one within-subjects factor (Recall Period) and two trial-level covariates (motion and signal variability) serving as explanatory variables. Random intercepts were estimated for each participant. Separate models were built for each seed region with the goal of identifying voxels whose correlations to a given seed exhibited a linear trend across the 3 Recall Periods (contrast weights: +1, 0, −1), which is mathematically equivalent to a direct contrast of the Today and 5-10 year ago conditions. Whole-brain correction for multiple comparisons was achieved for each map (*p* < .05) by setting a voxelwise *p* < .001 and a cluster extent of at least 18 voxels, as determined by 3dClustStim.

### Conjunction of connectivity clusters

The extent of overlap across connectivity difference maps was quantified using a conjunction image approach. Significant voxels/clusters in each map were binarized and subsequently summed. The resulting conjunction map therefore indicates, for each voxel, the number of discrete seed maps in which it appeared.

### Independent definition of scene-related cortical regions

The conjunction analysis suggested that a temporal gradient of connectivity was present among the univariate-defined parietal ROIs and regions of cortex likely associated with scene construction and scene perception. To independently define scene-selective cortex and test its correlations to posterior hippocampal ROIs, we used independent localizer data collected from 22 of the 40 subjects. We contrasted activity during blocks of scenes to that observed during all other task blocks and thresholded the resulting statistical map at a voxelwise *p* < .001 and a cluster extent of at least 18 voxels to achieve a whole-brain *p* < .05. Voxels exhibiting significantly greater activity for scenes than all other blocks were defined as scene-selective cortex. To ensure that no overlap was present between the scene-selective mask and the hippocampal regions within each subject, a conjunction of each participant’s left and right hippocampal masks was created and dilated by 1 voxel (using 3dmask_tool). The resulting voxels were excluded from the scene-selective mask to ensure that no overlapping voxels would influence the subsequent correlation analysis. The average correlation of voxels within this map to the left and right posterior hippocampal ROIs was computed and compared across conditions using the same LME modeling approach described for the voxelwise analysis, implemented in R using the lme4 package (84) (RRID: SCR_015654). Covariate-adjusted means were calculated for each condition with the R lsmeans package (85).

### Online Experiment 1

Participants were instructed to read event descriptions, rate each on the vividness of its description, and to identify the main location of the event in a short phrase. Participants read 3 event descriptions provided ratings using a 1-6 scale, where a score of 1 indicated “very generic” and a score of 6 indicated “very vivid”. Participants were further instructed that “A low score means the description is vague, generic, and lacking in detail. It may not even focus on a single event but instead wander from topic to topic.” They were also told that “A high rating means the event was described clearly and was full of detail, such as where an event occurred, who was there, and so on.” Participants were then given examples of low/high vividness descriptions.

After this instruction period, participants read each event description, provided a rating, and reported the main location for each event description. The experiment was self-paced, and participants were presented each description individually. The ordering of time periods was counterbalanced across participants. Participants were not told that event descriptions would come from different Recall Periods. A repeated measures ANOVA with the single factor of Recall Period was used to test for differences in ratings across the 3 temporal distances.

Event location responses served as our primary exclusion criteria. Participants who provided an incorrect response to any of the 3 events (either by not answering the question or providing an inaccurate response) were excluded.

### Online Experiment 2

Online Experiment 2 followed the procedure of Online Experiment 1, with the exception that participants now rated events on their overall level of detail, not vividness. In the updated instructions, participants were notified that a score of 1 indicated “very generic” and a score of 6 “very detailed”. All other aspects of this experiment were identical to Online Experiment 1.

## ACKNOWLEDGEMENTS

The authors thank Bess Bloomer and Gabrielle Reimann for assistance with data preparation; John Ingeholm and Bruce Prichard for technical assistance; Brendan Gaesser and Kevin Madore for sharing stimuli; and Michal Ramot, Andrew Persichetti, and Jason Avery for thoughtful discussions throughout this work. This work was supported by the NIMH Intramural Research Program (ZIAMH002920) and NIMH R01 MH060941 (to DLS).

## REFERENCES

1. E. Tulving, Elements of Episodic Memory (Oxford University Press, New York, 1983).

2. E. Tulving, Memory and consciousness. Canad. Psychol. 26, 1–12 (1985).

3. E. Svoboda, M. C. McKinnon, B. Levine, The functional neuroanatomy of autobiographical memory: A meta-analysis. Neuropsychologia 44, 2189–2208 (2006).

4. H. R. Hayama, K. L. Vilberg, M. D. Rugg, Overlap between the neural correlates of cued recall and source memory: Evidence for a generic recollection network. J Cog. Neurosci. 24, 1127–1137 (2012).

5. M. Boccia, A. Teghil, C. Guariglia, Looking into recent and remote past: Meta-analytic evidence for cortical re-organization of episodic autobiographical memories. Neurosci. Biobehav. Rev. 107, 84–95 (2019).

6. W. Scoville, B. Milner, Loss of recent memory after bilateral hippocampal lesions. J. Neurol. Neurosurg. Psychiatry 20, 11–21 (1957).

7. H. Eichenbaum, A cortical-hippocampal system for declarative memory. Nat. Rev. Neurosci. 1, 41–50 (2000).

8. L. R. Squire, D. L. Schacter, Eds., Neuropsychology of Memory (Guilford Press, New York, 2002), Third Ed.

9. T. Ribot, Diseases of memory (Kegan Paul, Trench, & Co., London, 1882).

10. W. R. Russell, P. W. Nathan, Traumatic amnesia. Brain 69, 280–300 (1946).

11. S. M. Zola-Morgan, L. R. Squire, The primate hippocampal formation: Evidence for a time-limited role in memory storage. Science 250, 288–290 (1990).

12. N. J. Broadbent, S. Gaskin, L. R. Squire, R. E. Clark, Object recognition memory and the rodent hippocampus. Learn. Mem. 17, 5–11 (2010).

13. P. Alvarez, L. R. Squire, Memory consolidation and the medial temporal lobe: A simple network model. Proc. Natl. Acad. Sci. U.S.A. 91, 7041–7045 (1994).

14. L. R. Squire, L. Genzel, J. T. Wixted, R. G. Morris, Memory consolidation. Cold Spring Harb Perspect Biol 7, a021766 (2015).

15. M. F. Carr, S. P. Jadhav, L. M. Frank, Hippocampal replay in the awake state: a potential substrate for memory consolidation and retrieval. Nat Neurosci 14, 147–153 (2011).

16. L. Nadel, M. Moscovitch, Memory consolidation, retrograde amnesia and the hippocampal complex. Curr. Opin. Neurobiol. 7, 217–227 (1997).

17. M. J. Sekeres, G. Winocur, M. Moscovitch, The hippocampus and related neocortical structures in memory transformation. Neurosci. Lett. 680, 39–53 (2018).

18. F. C. Bartlett, Remembering (Cambridge University Press, Cambridge, 1932).

19. A. D’Argembeau, M. Van der Linden, Phenomenal characteristics associated with projecting oneself back into the past and forward into the future: Influence of valence and temporal distance. Conscious. Cognit. 13, 844–858 (2004).

20. A. P. Yonelinas, C. Ranganath, A. D. Ekstrom, B. J. Wiltgen, A contexual binding theory of episodic memory: systems consolidation reconsidered. Nat. Rev. Neurosci. 20, 364–375 (2019).

21. P. Kundu et al., Integrated strategy for improving functional connectivity mapping using multiecho fMRI. Proc. Natl. Acad. Sci. U.S.A. 110, 16187–16192 (2013).

22. B. Gaesser, D. C. Sacchetti, D. R. Addis, D. L. Schacter, Characterizing age-related changes in remembering the past and imagining the future. Psychol. Aging 26, 80 (2011).

23. B. Levine, E. Svoboda, J. F. Hay, G. Winocur, M. Moscovitch, Aging and autobiographical memory: Dissociating episodic from semantic retrieval. Psychol. Aging 17, 677–689 (2002).

24. D. Cousineau, Confidence intervals in within-subject designs: A simpler solution to Loftus and Masson’s method Tutor Quant Methods Psychol 1, 42–45 (2005).

25. A. W. Gilmore et al., Dynamic content reactivation supports naturalistic autobiographical recall in humans. J. Neurosci. Advanced online access (2020).

26. J. Poppenk, H. R. Evensmoen, M. Moscovitch, L. Nadel, Long-axis specialization of the human hippocampus. Trends Cog. Sci. 17, 230–240 (2013).

27. I. K. Brunec et al., Multiple scales of representation along the hippocampal anteroposterior axis in humans. Curr. Biol. 28, 2129–2135.e2126 (2018).

28. D. S. Marcus et al., Informatics and data mining: Tools and strategies for the Human Connectome Project. Front. Neuroinform. 5, 4 (2011).

29. A. P. Yonelinas, L. J. Otten, K. N. Shaw, M. D. Rugg, Separating the brain regions involved in recollection and familiarity in recognition memory. J. Neurosci. 25, 3002–3008 (2005).

30. K. B. McDermott, K. K. Szpunar, S. E. Christ, Laboratory-based and autobiographical retrieval tasks differ substantially in their neural substrates. Neuropsychologia 47, 2290–2298 (2009).

31. H. Kim, Differential neural activity in the recognition of old versus new events: An activation likelihood estimation meta-analysis. Hum. Brain Mapp. 34, 814–836 (2013).

32. A. W. Gilmore, S. M. Nelson, K. B. McDermott, A parietal memory network revealed by multiple MRI methods. Trends Cog. Sci. 19, 534–543 (2015).

33. H.-Y. Chen, A. W. Gilmore, S. M. Nelson, K. B. McDermott, Are there multiple kinds of episodic memory? An fMRI investigation comparing autobiographical and recognition memory tasks. J. Neurosci. 37, 2764–2775 (2017).

34. J. D. Power, B. L. Schlaggar, S. E. Petersen, Recent progress and outstanding issues in motion correction in resting state fMRI. Neuroimage 105, 536–551 (2015).

35. S. J. Gotts, A. W. Gilmore, A. Martin, Brain networks, dimensionality, and global signal averaging in resting-state fMRI: Hierarchical network structure results in low-dimensional spatiotemporal dynamics. Neuroimage 25, 116289 (2020).

36. K. Jasmin et al., Overt socail interaction and resting state in young adult males with autism: Core and contextual neural features. Brain 142, 808–822 (2019).

37. R. G. Benoit, D. L. Schacter, Specifying the core network supporting episodic simulation and episodic memory by activation likelihood estimation. Neuropsychologia 75, 450–457 (2015).

38. A. W. Gilmore, S. M. Nelson, K. B. McDermott, The contextual association network activates more for remembered than for imagined events. Cereb. Cortex 26, 611–617 (2016).

39. A. W. Gilmore, S. M. Nelson, H.-Y. Chen, K. B. McDermott, Task-related and resting-state fMRI identify distinct networks that preferentially support remembering the past and imagining the future. Neuropsychologia 110, 180–189 (2018).

40. D. Hassabis, D. Kumaran, E. A. Maguire, Using imagination to understand the neural basis of episodic memory. J. Neurosci. 27, 14365–14374 (2007).

41. C. Baldassano, A. Esteva, L. Fei-Fei, D. M. Beck, Two distinct scene-processing networks connecting vision and memory. eNeuro 3, ENEURO–0178 (2016).

42. S. L. Mullally, E. A. Maguire, Memory, imagination, and predicting the future: A common brain mechanism? The Neuroscientist 20, 220–234 (2014).

43. D. C. Rubin, S. Umanath, Event memory: A theory of memory for laboratory, autobiographical, and fictional events. Psychol. Rev. 122, 1–23 (2015).

44. J. Robin, Spatial scaffold effects in event memory and imagination. WIREs Cognitive Science 23, e1462 (2018).

45. H. J. Spiers, E. A. Maguire, Thoughts, behavior, and brain dynamics during naviation i the real world. Neuroimage 31, 1826–1840 (2006).

46. P. V. Rekkas, R. T. Constable, Evidence that autobiographic memory retrieval does not become independent of the hippocampus: An fMRI study contrasting very recent with remote events. J Cog. Neurosci. 17, 1950–1961 (2005).

47. D. R. Addis, D. L. Schacter, Constructive episodic simulation: Temporal distance and detail of past and future events modulate hippocampal engagement. Hippocampus 18, 227–237 (2008).

48. A. Martin, C. L. Wiggs, J. Weisberg, Modulation of human medial temporal lobe activity by form, meaning, and experience. Hippocampus 7, 587–593 (1997).

49. A. Viard et al., Hippocampal activation for autobiographical memories over the entire lifetime in healthy aged subjects: An fMRI study. Cereb. Cortex 17, 2453–2467 (2007).

50. K. P. Madore, B. Gaesser, D. L. Schacter, Constructive episodic simulation: Dissociable effects of a specificity induction on remembering, imaging, and describing in younger and older adults. J Exp Psychol -Learn Mem Cogn 40, 609–622 (2014).

51. A. Gilboa, G. Winocur, C. L. Grady, S. J. Hevenor, M. Moscovitch, Remembering our past: functional neuroanatomy of recollection of recent and very remote personal events. Cereb. Cortex 14, 1214–1225 (2004).

52. A. D. Wagner, B. J. Shannon, I. Kahn, R. L. Buckner, Parietal lobe contributions to episodic memory retrieval. Trends Cog. Sci. 9, 445–453 (2005).

53. M. L. Rosen, C. E. Stern, K. J. Devaney, D. C. Somers, Cortical and subcortical cotributions to long-term memory-guided visuospatial attention. Cereb. Cortex 28, 2935–2947 (2018).

54. A. W. Gilmore, S. E. Kalinowski, S. C. Milleville, S. J. Gotts, A. Martin, Identifying task-general effects of stimulus familiarity in the parietal memory network. Neuropsychologia 124, 31–43 (2019).

55. S. M. Nelson, K. M. Arnold, A. W. Gilmore, K. B. McDermott, Neural signatures of test-potentiated learning in parietal cortex. J. Neurosci. 33, 11754–11762 (2013).

56. K. K. Szpunar, P. L. St Jacques, C. A. Robbins, G. S. Wig, D. L. Schacter, Repetition-related reductions in neural activity reveal component processes of mental simulation. Social Cognitive and Affective Neuroscience 10.1093/scan/nst035 (2014).

57. K. K. Szpunar, H. G. Jing, R. G. Benoit, D. L. Schacter, Repetition-related reductions in neural activity during emotional simulations of future events. PLoS ONE 10, e0138354 (2015).

58. A. L. Devitt, P. P. Thakral, K. K. Szpunar, D. R. Addis, D. L. Schacter, Age-related changes in repetition suppression of neural activity during emotional future simulation. Neurobiology of Aging 94, 287–297 (2020).

59. A. Takashima et al., Declarative memory consolidation in humans: A prospective functional magnetic resonance imaging study. Proc. Natl. Acad. Sci. U.S.A. 103, 756–761 (2006).

60. H. M. Bonnici, E. A. Maguire, Detecting representations of recent and remote autobiographical memories in vmPFC and hippocampus. J. Neurosci. 32, 16982–16991 (2012).

61. H. M. Bonnici, E. A. Maguire, Two years later – Revisiting autobiographical memory representations in vmPFC and hippocampus. Neuropsychologia 110, 159–169 (2018).

62. D. N. Barry, M. J. Chadwick, E. A. Maguire, Nonmonotonic recruitment of ventromedial prefrontal cortex during remote memory recall. PLoS biology 16, e2005479 (2018).

63. D. N. Barry, E. A. Maguire, Remote memory and the hippocampus: A constructive critique. Trends Cog. Sci. 23, 128–142 (2019).

64. C. McCormick, D. N. Barry, A. Jafarian, G. R. Barnes, E. A. Maguire, vmPFC drives hippocampal processing during autobiographical memory recall regardless of remoteness. Cereb. Cortex 30, 5972–5987 (2020).

65. M. A. Ferguson et al., A human memory circuit derived from brain lesions causing amnesia. Nature Communications 10, 1–9 (2019).

66. W. D. Stevens, M. H. Tessler, C. S. Peng, A. Martin, Functional connectivity constrains the category-related organization of the human ventral occipitotemporal cortex. Hum. Brain Mapp. 36, 2187–2206 (2015).

67. J. W. Peirce, PsychoPy -Psychophysics software in python. J. Neurosci. Meth. 162, 8–13 (2007).

68. E. M. Gordon et al., Precision functional mapping of individual human brains. Neuron 95, 791–807.e797 (2017).

69. J. Yuan, M. Liberman, Speaker identification on the SCOTUS corpus. Proceedings of Acoustics ‘08 (2008).

70. P. Boersma, V. van Heuven, Praat, a system for doing phonetics by computer. Glot International 5, 341–345 (2001).

71. A. Martin, GRAPES—Grounding representations in action, perception, and emotion systems: How object properties and categories are represented in the human brain. Psychon Bull Rev 23, 979–990 (2016).

72. Y. Benjamini, Y. Hochberg, Controlling the false discovery rate: A practical and powerful approach to multiple testing. J R Stat Soc Series B Stat Methodol 51, 289–300 (1995).

73. R. W. Cox, AFNI: software for analysis and visualization of functional magnetic resonance images. Comput Biomed Res 29, 162–173 (1996).

74. P. Kundu, S. J. Inati, J. W. Evans, W.-M. Luh, P. A. Bandettini, Differentiating BOLD and non-BOLD signals in fMRI time series using multi-echo EPI. Neuroimage 60, 1759–1770 (2012).

75. P. Kundu et al., Multi-echo fMRI: A review of applications in fMRI denoising and analysis of BOLD signals. Neuroimage 154, 59–80 (2017).

76. J. D. Power et al., Ridding fMRI data of motion-related influences: Removal of signals withdistinct spatial and physical bases in multiecho data. Proc. Natl. Acad. Sci. U.S.A. 115, E2105–E2114 (2018).

77. D. I. Donaldson, S. E. Petersen, R. L. Buckner, Dissociating memory retrieval processes using fMRI: evidence that priming does not support recognition memory. Neuron 31, 1047–1059 (2001).

78. K. M. Visscher et al., Mixed blocked/event-related designs separate transient and sustained activity in fMRI. Neuroimage 19, 1694–1708 (2003).

79. S. E. Petersen, J. W. Dubis, The mixed block/event-related design. Neuroimage 62, 1177–1184 (2012).

80. R. W. Cox, G. Chen, D. R. Glen, R. C. Reynolds, P. A. Taylor, fMRI clustering and false-positive rates. Proc. Natl. Acad. Sci. U.S.A. 114, E3370–E3371 (2017).

81. R. W. Cox, G. Chen, D. R. Glen, R. C. Reynolds, P. A. Taylor, fMRI clustering in AFNI: False-positive rates redux. Brain Connect 7, 152–171 (2017).

82. A. Eklund, T. E. Nichols, H. Knutsson, Cluster failure: Why fMRI inferences for spatial extent have inflated false-positive rates. Proc. Natl. Acad. Sci. U.S.A. 113, 7900–7905 (2016).

83. G. Chen, Z. S. Saad, J. C. Britton, D. S. Pine, R. W. Cox, Linear mixed-effects modeling approach to FMRI group analysis. Neuroimage 73, 176–190 (2013).

84. D. Bates, M. Mächler, B. Bolker, S. Walker, Fitting linear mixed-effects models using lme4. J Stat Softw 61, 1–48 (2015).

85. R. Lenth, Least-squares means: The R package lsmeans. J Stat Softw 69, 1–33 (2016).

