## Supplementary Figures and Tables for "The human hippocampus plays a time-limited role in retrieving autobiographical memories"

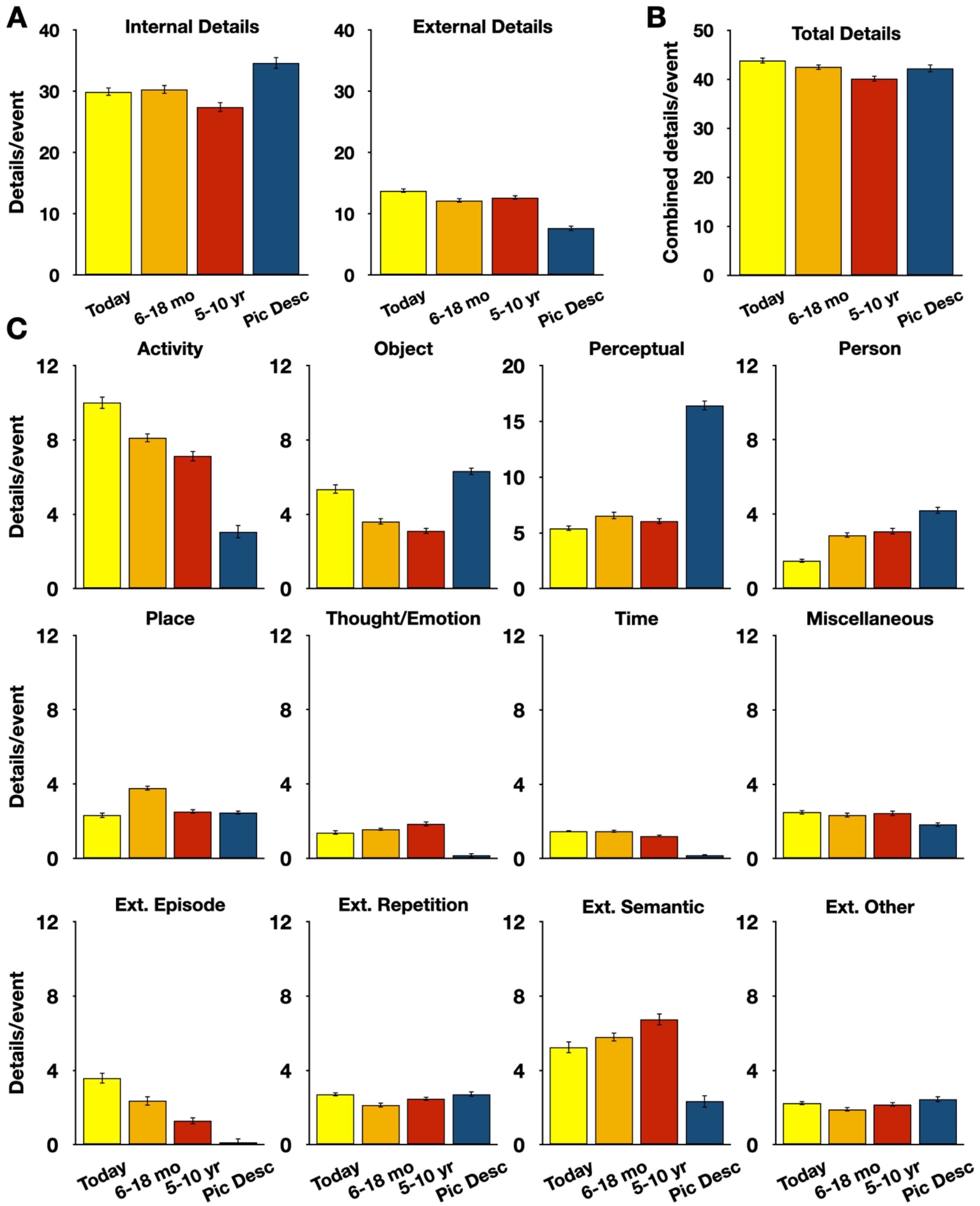


**Figure S1.** Breakdown of Internal and External details across all task conditions, including the Picture Description baseline. A) Average counts of Internal and External event categories. B) Average combined totals (Internal + External) for each task condition. C) Average counts of detail subtypes for each task condition. Error bars denote within-subject standard error.

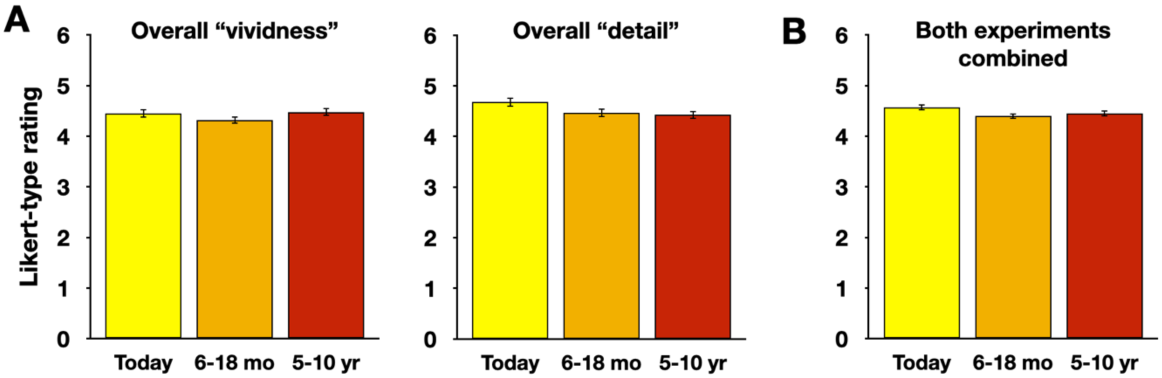


**Figure S2.** Subjective ratings of overall vividness and detail from two Online Experiments. A) Neither ratings of "vividness"(N = 129) nor of "detail"(N = 144) differed across Recall Periods, suggesting that qualitative differences were not observed across recent and remote events in the current dataset. B) Combining data into a single large group likewise did not result in a significant effect of Temporal Distance on subjective event description ratings. Error bars denote within-subject standard error.

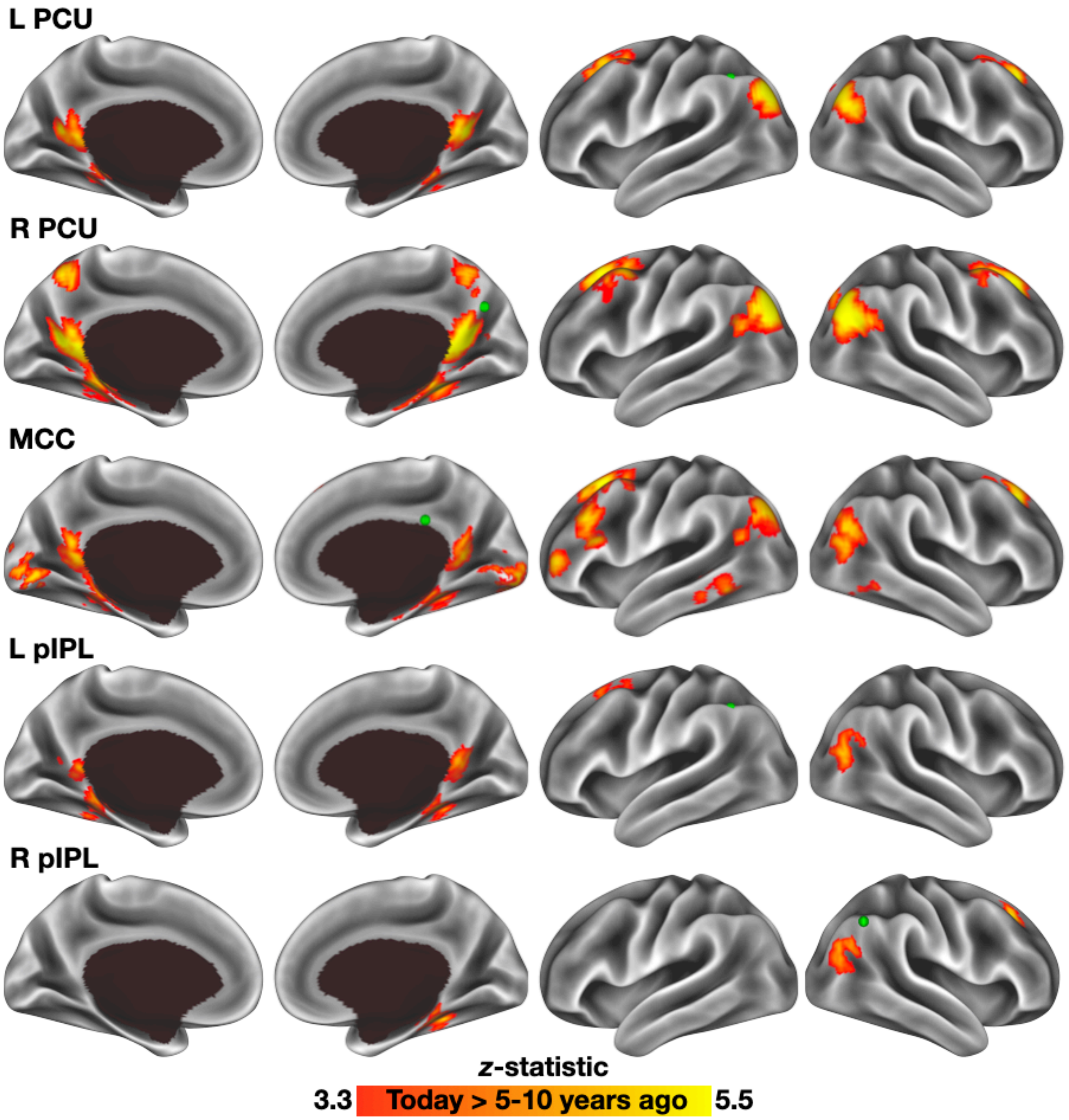


**Figure S3.** Whole brain contrast results for ROIs that exhibited significant differences in connectivity between the Today and 5-10 year ago conditions. Green nodes depict the center of each seed region. Warm colors represent greater connectivity during the recall of recent than of remote events.

Table S1. List of Internal and External detail categories.

| **Category** | **Description** |
|---|---|
| ***Internal*** | *Details associated with a spatially and temporally specific event* |
| Activity | Something done or undertaken by a person or object |
| Object | A non-living entity |
| Perceptual Detail | Sensory details, including relative spatial positions and durations |
| Person | A human entity |
| Place | Description of the "where" of an event |
| Thought/Emotion | Experienced and attributed emotional states, descriptions of ongoing thoughts |
| Time | A description of "when" an event took place, including time of day, season, etc. |
| Miscellaneous | Contains details that did not occur frequently enough for separate modeling, includes descriptions of animals, body parts, weather phenomena, etc. |
| ***External*** | *Details not specific to the reported event* |
| Episodic | Spatially and temporally specific details from occurrences other than the main event being described |
| Repetition | Repetitions of "Internal" details |
| Semantic | General knowledge or background, often either tangential to or offered in support of the described event |
| Other | Editorial comments, banter, etc. |
